# Cell-Free Protein Expression in Polymer Materials

**DOI:** 10.1101/2023.08.18.553881

**Authors:** Marilyn S. Lee, Jennifer A. Lee, John R. Biondo, Jeffrey E. Lux, Rebecca M. Raig, Pierce N. Berger, Casey B. Bernhards, Danielle L. Kuhn, Maneesh K. Gupta, Matthew W. Lux

## Abstract

While synthetic biology has advanced complex capabilities like sensing and molecule synthesis in aqueous solutions, important applications may also be pursued for biological systems in solid materials. Harsh processing conditions used to produce many synthetic materials such as plastics makes incorporation of biological functionality challenging. One technology that shows promise in circumventing these issues is cell-free protein synthesis (CFPS), where core cellular functionality is reconstituted outside the cell. CFPS enables genetic functions to be implemented without the complications of membrane transport or concerns over cellular viability or release of genetically modified organisms. Here we demonstrate that dried CFPS reactions have remarkable tolerance to heat and organic solvent exposure during the casting processes for polymer materials. We demonstrate the utility of this observation by creating plastics that have spatially patterned genetic functionality, produce antimicrobials *in situ*, and perform sensing reactions. The resulting materials unlock the potential to deliver DNA-programmable bio-functionality in a ubiquitous class of synthetic materials.

## 1. INTRODUCTION

Nature abounds with materials capable of complex dynamic functions that conventional material science cannot match. From tissues like muscles and skin to hard shells and bones, these bio-composites have multiple functions, can change over time, and respond to many stimuli. Even more remarkably, the blueprints for the synthesis, construction, and function of these materials are encoded in the DNA of the organisms that create them. Synthetic biology enables the design of similarly capable composites by applying rapidly developing tools to engineer organisms and use them to functionalize materials.^1-5^ Conceptually, biologically-enabled materials can sense damage or threats and provide outputs like color change, fluorescence, or even electrical signal. Metabolic activity can produce useful molecules like therapeutics or antimicrobials *in situ*. Self-cleaning, self-repair, or self-degradation can be mediated by enzymes. All these functions are achieved by re-programming the DNA instructions that encode the desired properties.

A critical aspect of engineering such living materials is ensuring the viability of the organisms responsible for the intended biological functions. As a result, advances have focused on environments that are generally friendly to microbes, such as hydrogels^6-10^ or biofilms,^11-15^ or use organisms naturally adapted to a target environment, such as architectural materials, soil, seawater, or the human gut.^16-22^ Bacterial spores are sometimes used to enable the cells to withstand processing conditions or to extend the duration during which a living material can function.^8,23,24^ Often, it is not clear how to make nutrients available to microbes residing inside a material. Most approaches described by recent works involve soaking the object in a nutrient bath, which may not be appropriate for all applications and could limit the lifetime of the programmed functionality to feeding events.^23,25^ As a result, materials that are unfavorable to biological systems, whether in their processing or final form, remain off-limits for engineered living materials.

Plastics represent an important class of synthetic materials for which the processing is generally inhospitable to biological systems. Many categories of polymer materials exist with a wide range of synthesis methods, most of which involve conditions that are harsh for biological systems due to high temperatures, exposure to solvents, radical generation, or UV radiation. Some studies have shown the possibility of proteins surviving these conditions, whether by protecting the protein with additives or encapsulation,^26-29^ or engineering the enzyme to be more robust.^30-34^ While promising, these approaches limit the biological function to a single enzyme or set of enzymes without programmable dynamic response and require re-engineering the formulation or protein sequence variant for each new application.

One avenue that bypasses the drawbacks of both living cells and isolated enzymes to deliver programmable biological function to materials is cell-free protein synthesis (CFPS), where the basic cellular functions of transcription, translation, and metabolism are recapitulated outside the confines of the cell. CFPS systems offer an alternative to living cells for implementing bio-functionality in materials and can circumvent problems like genetic instability and concerns over environmental release of genetically modified organisms.^35-37^ CFPS reactions are complex mixtures of crude lysates or purified cellular components and excipient resources that enact DNA-encoded functions like sensing and production of proteins such as enzyme catalysts or therapeutics.^38-42^ CFPS reactions achieve these functions without the need to maintain cell viability and replication, and the lack of cell membrane enables dramatically more control over the reaction environment. Resources supplied to the reaction typically include nucleosides, amino acids, phosphorylated or sugar energy sources, buffer, and cofactors, with many different formulations reported to achieve different features and functions.^43-45^ Remarkable stability has been observed for freeze-dried CFPS powder stored at room temperature with desiccant for up to a year.^46-49^ Further, several commonly used dry formulations can tolerate exposure to a variety of organic solvents, although stability following solvent exposure can vary greatly depending on the individual components.^43^ Other work has shown that lysates enriched with specific proteins like Cas12, used in a sensing application, require lyo-protectants to maintain functionality.^50^ While there are many considerations when developing and optimizing CFPS systems to embed in synthetic materials, these reactions are open systems that allow unfettered access to the reaction environment, breaking the limitations of living systems.

Porous materials like fabrics, paper, and hydrogels have been functionalized with CFPS.^51-55^ In this study, we seek to establish a proof-of-concept for embedding CFPS reactions in solvent- and heat-cast polymer materials (Figure 1A). We test the limits of solvent- and heat-casting methods for dried CFPS powder in poly lactic-co-glycolic acid (PLGA) and polycaprolactone (PCL), respectively. Protein synthesis activity was recovered from films with embedded lyophilized CFPS powder after each of these casting methods. PLGA and PCL have differing characteristics that impact the performance of embedded CFPS, particularly with respect to water infiltration of the material. Several functions are demonstrated in solvent-cast CFPS-PLGA: constitutive production of a single fluorescent protein, patterned localized production of two fluorescent proteins in the same material, production of an antimicrobial colicin to inhibit *Escherichia coli* growth, and a toehold switch sensor responding to an RNA trigger. We conclude that polymer-embedded CFPS reactions are a promising new way to deliver complex, DNA-programmable biological activity in materials. We then discuss the needs for further developments and the new directions that polymers could take CFPS-driven synthetic biology.

**Figure 1.**
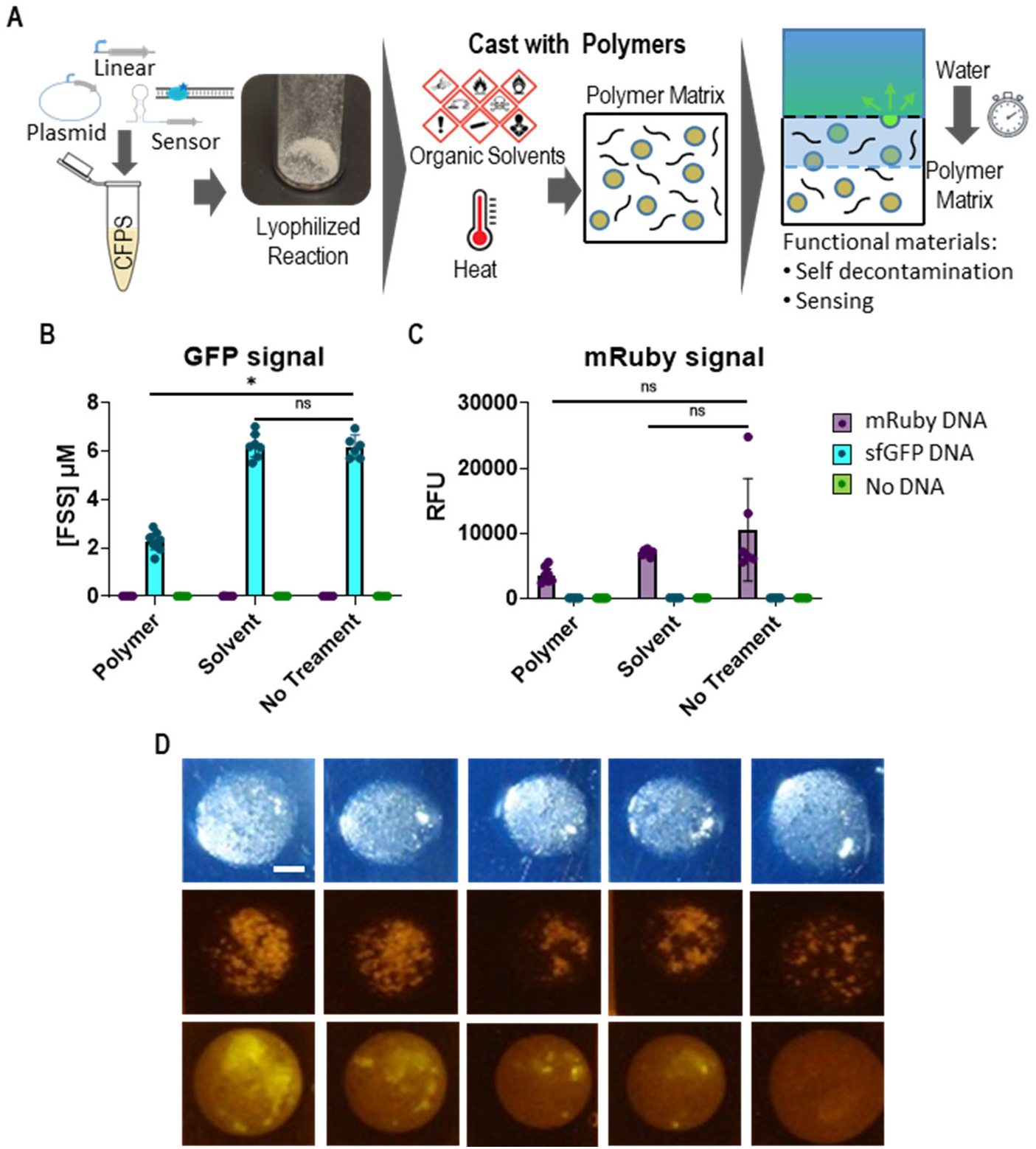
CFPS powder cast into polymers. (A) Graphic describing the incorporation of CFPS reactions in a polymer matrix: reaction assembly with many possible DNA configurations dictating function and lyophilization to form a dry CFPS powder (left), polymer casting via harsh solvent or heat treatments to embed the stable CFPS powder within a non-aqueous polymer matrix (center), and reactivation via water permeation into the CF-polymer composite and/or hydrolysis of the polymer (right). (B) Endpoint sfGFP fluorescence signal in units of equivalent fluorescein concentration ([FSS]) for samples containing mRuby DNA, sfGFP DNA or no DNA. CFPS powder samples were mixed with PLGA and acetone, acetone alone, or untreated. The mean and individual 8 h endpoints are shown, n≥ 6; error bars are the 95% confidence interval. (C) mRuby fluorescence in relative fluorescence units (RFU) for the same samples described in (B). (D) Representative photographs of five solvent-cast CFPS-PLGA films containing plasmid encoding sfGFP before (top and middle) and 3 h after (bottom) rehydration. Top image was taken under white light. Middle and bottom images taken under blue light with orange filter to capture sfGFP fluorescence. Scale bar is 2 mm.

## 2. RESULTS

### Solvent-cast CFPS-PLGA

Dried CFPS powders derived from several different reaction recipes are stable to room temperature storage and organic solvent exposure.^46-49^ We sought to harness this ability to test encapsulation of CFPS within solvent-cast polymer. The CFPS reaction formulation utilized in this study is comprised of *E. coli* lysate mixed with the PANOx-SP solution of protein synthesis resources, cofactors, and buffer.^56^ PLGA was the first polymer tested because it is biocompatible, can readily absorb water, and undergoes gradual hydrolysis of its polymer chains to release encapsulated cargo. These properties indicated it might give favorable conditions to reactivate the CFPS activity upon hydration. Acetone is suitable to dissolve PLGA to cast films and is one of the solvents previously determined to be compatible with CFPS activity for this formulation.^46^

As a first test, the recovery of protein synthesis activity was measured after dry reaction components were exposed to a solution of polymer in acetone. CFPS reactions containing plasmid DNA encoding a constitutive T7 promoter driving expression of either sfGFP or mRuby fluorescent proteins were deposited in a solvent resistant microplate and lyophilized. Then, samples were exposed to either acetone alone, PLGA dissolved in acetone, or left untreated. Acetone was allowed to evaporate from the wells, leaving behind dry polymer and CFPS components. Each well was then rehydrated with water and the production of fluorescence was monitored during incubation. The results for mRuby and sfGFP products bore similar trends (Figure 1B, C, Figure S1). Acetone exposure did not show any decrease in productivity, as expected. The addition of polymer caused endpoint mean productivity to drop to 37% of untreated mean levels for sfGFP, p<<0.05. The difference in mean for mRuby signal was not significant between treatments, though this may be due to high experimental variability in the no treatment control case. Though activity dropped for at least one reporter, this result verified that CFPS can work when processed with solvent-cast polymers and encouraged further exploration of polymer casting.

To proceed to solvent-casting films, it was necessary to form a suspension of CFPS powder in acetone-PLGA solution and transfer the mixture to cast on a glass substrate. We found that preparation methods of the lyophilized CFPS powder were important to achieve consistent performance and shelf stability up to 10 weeks (Supplementary Information, Supplementary Table S1). CFPS powder was combined with acetone-PLGA solution to form a suspension. The suspension was drop-cast onto a glass surface and the acetone was allowed to evaporate to form a CFPS-PLGA film (Figure 1D). Films were submerged in water and incubated at 37°C to observe the appearance of protein product. sfGFP fluorescence was observed via photographs under blue light and was visible by eye, demonstrating CFPS functionality in the solvent-cast films. To our knowledge, this is the first such demonstration of complex metabolic reactions and protein synthesis function from a non-living system in a non-hydrogel material. In these initial experiments, sfGFP fluorescence was not uniformly distributed in all films, with some films failing to produce fluorescence, indicating non-uniform mixing and dispersion of the CFPS reagents in the polymer and solvent suspension. Nonetheless, this proof-of-concept encouraged further development to achieve more reliable and more complex DNA-programmed functionality in CFPS-polymer composites.

### Heat-cast CFPS-PCL

High-heat processing is another major way to form solid objects from polymer materials. Previous studies examining heat tolerance of dried CFPS reagents focused on long-term exposure over many days or weeks to temperatures ranging from 20–50°C to demonstrate shelf stability.^47,49,57^ We pursued heat-casting experiments for CFPS-polymer composites to learn how a polymer matrix might affect the heat tolerance of CFPS reagents. Many polymers melt at temperatures much higher than 100°C, but there are some that melt at lower temperatures better aligned with the stability limitations for CFPS, including PCL with a melting point of 60°C. PCL is also biocompatible and undergoes slow hydrolytic degradation, potentially lending itself to gradual rehydration and reactivation of CFPS components over time. These properties make PCL a good candidate for initial tests.

CFPS powder containing plasmid DNA expressing mRuby fluorescent protein was embedded within heat-cast PCL films. Pure PCL films of about 200-micron thickness were formed by pressing a PCL pellet between plates heated to 80°C using a manual heat press (Figure 2A, B top). Aqueous CFPS reactions were dropped on PCL bottom films and lyophilized in place. Then a second film was placed on top to form a sandwich around the dry CFPS reagents, and the stack was re-pressed for 5 seconds to embed the powder in heated polymer (Figure 2B, bottom). Intact or cut PCL films with embedded CFPS were rehydrated with water and incubated to monitor mRuby production via microscopy (Figure 2C, D). Increasing fluorescence levels over time were measured in films that contained DNA but were not detected in films where DNA was left out of the CFPS mixture. Increased fluorescence was observed localized at cut edges compared to intact films. Overall, these results are the first to show that it is possible to recover CFPS activity within a polymer material after a heat-pressing process. No adjustments to CFPS reaction formulation have yet been explored for PCL casting, and further improvements might be achieved by using lyo-stabilizers like trehalose or maltodextrin which have been demonstrated to improve CFPS heat stability.^57^

**Figure 2.**
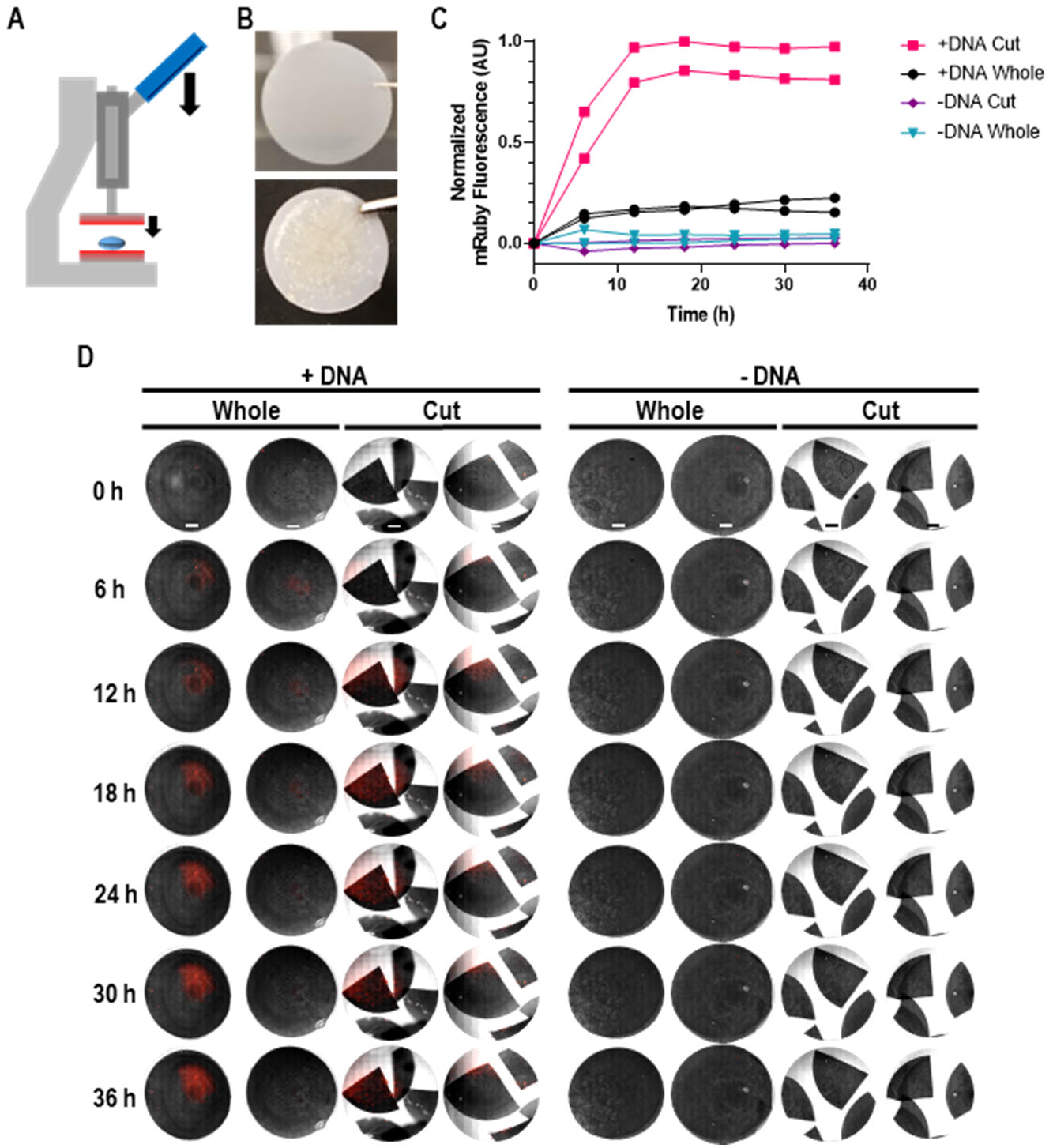
Heat-cast CFPS-PCL. (A) Diagram of heat press used to form PCL films, (B) Photographs of PCL film without (top) and with (bottom) embedded CFPS powder, (C) Quantified fluorescence from microscopy imaging of hydrated PCL films with and without mRuby plasmid DNA, either whole or cut, (D) Microscopy images of the PCL films quantified in (C), scale bars are 2 mm.

While CFPS-PLGA films exhibited clear water intrusion via clouding and swelling of the films and produced fluorescent protein across the entire film, whole CFPS-PCL films did not produce as much fluorescence as cut PCL films. Interestingly, dye intrusion experiments indicate water infiltrates into cut CFPS-PCL composite, though no intrusion was observed with cut pure PCL (Figure S2). This indicates that the embedded CFPS material can cause water-accessible voids and porosity in the otherwise hydrophobic PCL. Pure PCL is much more hydrophobic than PLGA, preventing water intrusion on the time scales of our experiments. Due to our sandwich heat-pressing method, CFPS material is laminated between PCL layers and is likely less exposed to water in whole films verses cut. Possible alternate explanations include oxygen limitations on CFPS or protein maturation within the PCL or decreased optical impedance by the polymer leading to better fluorescence signal at the edge. While this data does not conclusively pinpoint the cause of localized edge activity in PCL, the difference in activity trends and morphology between PCL and PLGA polymers highlights that polymers with different properties could be selected to tune the timing, location, and degree of CFPS activation.

### Localized function in CFPS-PLGA

A hallmark of natural and engineered biological materials is spatial patterning, including DNA-encoded sense and respond functions at the micro scale that leads to emergent macroscale properties.^58,59^ The ability to embed CFPS biological activity offers the opportunity to craft spatial localization of functionalities within a polymer material. Patterning of multiple functions within the same contiguous material offers interesting properties like controlled diffusion that can influence the interplay and behaviors of reactions in the material.^7,60-63^ Towards this concept, the process to solvent-cast CFPS reagents into films was improved and two different PLGA-CFPS “inks” suspended in acetone solvent were created. One ink contained plasmid DNA encoding mRuby, the other encoding sfGFP; a third had no DNA as a control. Uniformity of inks and the resulting films was improved by using a pestle to finely crush the CFPS powder in the presence of the acetone-PLGA solution, forming a more stable suspension (Figure 3A). Then, fluorescent protein production in solvent-cast films was more sensitively monitored via an inverted microscope.

**Figure 3.**
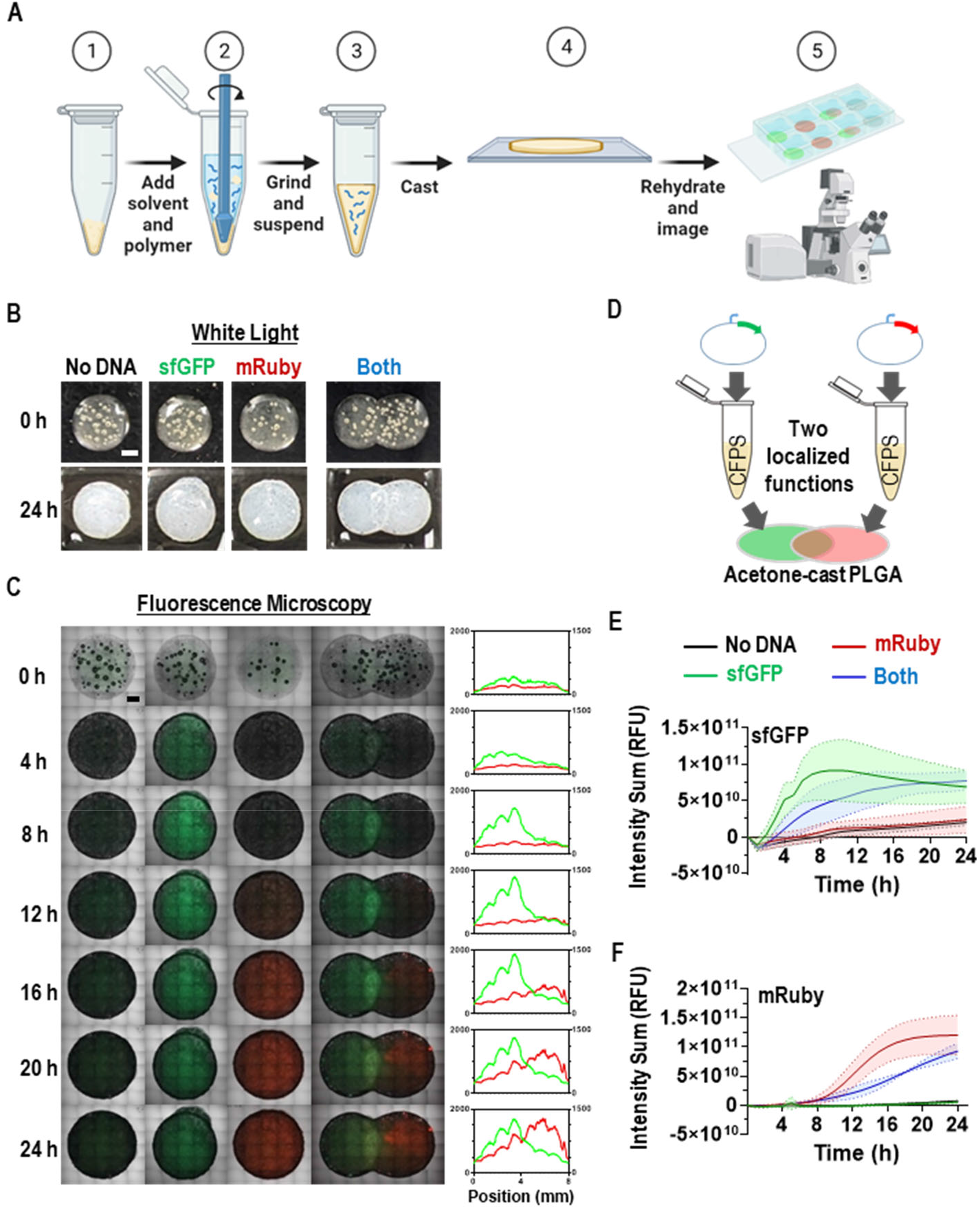
Microscopy of localized functions in CFPS-PLGA films. (A) Sequence of steps for improved solvent-cast CFPS-PLGA process, (B) Representative white light photographs of CFPS-PLGA films before rehydration and after 24 h incubation with water. Scale bar is 1 mm. Additional replicates are depicted in Figure S4. (C) Representative tiled microscopy images of whole CFPS-PLGA films. DIC, sfGFP (green), and mRuby (red) fluorescence channels are merged. Scale bar is 1 mm. Additional replicates and unmerged images are depicted in Figures S5 and S6. Quantitative fluorescence intensity profiles across the dual color film for green and red channels is shown at right. Left y-axis is green fluorescence and right y-axis is red fluorescence, units are RFU. (D) Diagram of two CFPS-PLGA “inks” functionalized with different DNA cast into adjacent and overlapping films. Quantified fluorescence intensity in whole CFPS-PLGA films measured by microscope for sfGFP (E) and mRuby (F) channels. The samples measured were films with no DNA (black), sfGFP DNA (green), mRuby DNA (red), or overlapping films containing sfGFP and mRuby DNA (blue). Error bars represent standard deviation for fluorescence measured in n=3 films.

Differential interference contrast (DIC) was used to capture the morphology of the CFPS-PLGA film compared to PLGA alone (Figure S3). The embedded CFPS powder particles are irregular and vary greatly in size. PLGA film is initially transparent; addition of water causes an increase of opacity due to hydrolysis of the PLGA (Figures 3B, S4). The appearance of fluorescence in green and red widefield channels is observed for each film, using time course and image stitching features to capture activity over the area of an entire film (Figures 3C, S5, S6, Supplementary videos S1–4).

For two-color films, a droplet of sfGFP ink was deposited on glass and allowed to dry to form a film. The mRuby ink was then deposited in contact with the first film and dried. The result was a combined PLGA film with two differently functionalized regions (Figure 3D). Fluorescence intensity was quantified for all film configurations (Figure 3E, F). As expected, the films without DNA do not show increased levels of fluorescence over time, while sfGFP and mRuby films produce green and red fluorescence, respectively. The mRuby signal typically takes much longer to mature compared to sfGFP signal in CFPS reactions, and this trend is apparent for CFPS-films as well. Comparing the dynamics, sfGFP fluorescence in CFPS films takes nearly 10 hours to reach maximum signal whereas it takes around 4 h for CFPS solutions (Figure S7). This difference in rate is expected given a much longer maturation time for mRuby compared to GFP.^64,65^ Visual inspection of the two-color films indicates localization of green and red fluorescence in the expected distribution, with more green on the left and more red on the right. Intensity profiles highlight the fluorescence distribution. Some protein or DNA likely diffuse between films since there is a small amount of mRuby signal distributed across the sfGFP DNA film, and vice versa (Figures 3C, S5, S6). Total integrated fluorescence values for each film over time indicate overall mRuby and sfGFP productivity in the two-color films is similar to the single-color films (Figure 3E–F).

### Production of colicin in CFPS-PLGA

We next sought to demonstrate the capacity of CFPS to deliver a biological payload not typically possible with polymer materials by expressing an antimicrobial protein, colicin E1, in PLGA films. Embedding the production of colicin could yield self-decontaminating materials that may be useful to prevent colonization of harmful bacteria in devices like medical implants, ventilators, food processing equipment, or personal protective equipment. Antimicrobial proteins like colicins are attractive alternatives to traditional antibiotics due to highly specific activity, huge diversity in nature, and extant tools to explore the mutational space to increase diversity even further. However, they are typically sensitive to proteases and other modes of degradation, and each type of antimicrobial protein has differing degrees of heat or shelf stability.^66-68^ Moreover, colicins can be difficult to produce via cell-based biomanufacturing because of their toxicity to live bacterial hosts.^55,69,70^ Synthesizing these molecules *in situ* using non-living CFPS sidesteps many of the limitations of antimicrobial proteins and potentially enables their delivery in materials. Because these materials are reprogrammable by changing the DNA sequence, the approach could also reduce the effort required to deliver a different antimicrobial activity without having to tailor the process to the requirements of each individual protein.

Linear PCR product encoding colicin under the control of a T7 promoter was added to the CFPS solution (Figure 4A). Using linear DNA eliminates the need to construct plasmid DNA in a live host, avoiding toxicity issues caused by leaky expression of the potent antimicrobial protein. Creating challenging plasmids can take weeks or months, if it is successful at all; the linear DNA approach only requires a few days of waiting for the delivery of commercially synthesized DNA. GamS nuclease inhibitor was also added to protect the linear template from degradation in the CFPS reaction. After lyophilization, the CFPS reaction material with or without colicin DNA was rehydrated in an incubator after no treatment, treatment with pure acetone, or treatment with PLGA dissolved in acetone. Samples of the CFPS or CFPS-polymer supernatant solutions were then added to *E. coli* K-12 subcultures to assess growth inhibition activity by colicin. All CFPS reactions containing the colicin DNA exerted significant growth inhibition on the *E. coli* cultures relative to no DNA controls (Figure 4B). Growth inhibition suggests active colicin was produced in solution even after exposure to solvent and polymer casting conditions. This result demonstrates the potential of delivering useful proteins *in situ* simply by including the requisite DNA.

**Figure 4.**
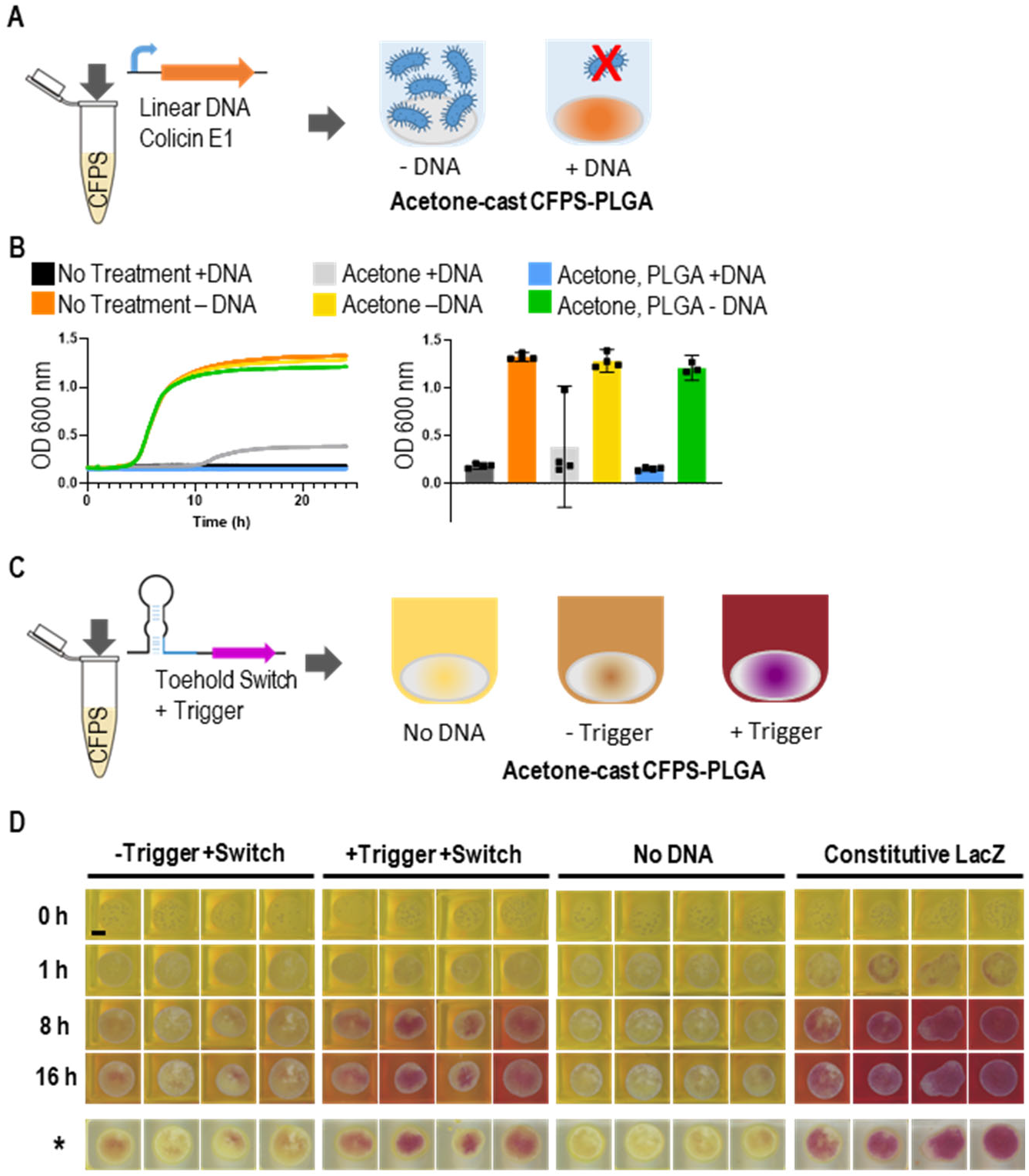
Expanded functionality in solvent-cast polymer. (A) Diagram of antimicrobial polymer film capable of preventing *E. coli* growth formed by embedding CFPS functionalized with linear DNA encoding colicin E1 in acetone-cast CFPS-PLGA, (B) Optical density growth time course (left) and endpoint (right) data are shown for *E. coli* K-12 cultures exposed to CFPS reaction solutions with or without colicin DNA that were rehydrated in an incubator after no treatment, treatment with pure acetone, or treatment with PLGA dissolved in acetone. Error bars represent the 95% confidence interval, n≥3, (C) A polymer film sensor was formed by including a plasmid encoding a LacZ enzyme under control of a toehold RNA switch as well as a plasmid encoding the cognate trigger RNA in polymer-CFPS, (D) Images of CPRG color change in CFPS-PLGA reactions containing the toehold switch and trigger plasmid DNA, the switch plasmid without trigger, constitutively expressed LacZ positive control plasmid DNA, or no DNA. Four time points are shown with films submerged in CPRG solution, the final timepoint is also depicted with CPRG solution removed (*) to show the pigmented films. Scale bar is 2 mm.

### Sensor functionality in CFPS-PLGA

Finally, we set out to demonstrate a dynamic sense-and-respond function, another universal capability of biology not generally found in polymer materials. We selected an RNA toehold switch sensor, Switch H,^48^ as representative of a class of sensors that have been used to detect many pathogens and disease biomarkers. By linking the sensor to a colorimetric reporter enzyme, plastics that change color in the presence of a pathogen or other analyte of interest become possible. Lyophilized CFPS powder samples were prepared that contained combinations of two plasmids, one encoding LacZ (β-galactosidase) under the translational control of a toehold switch, and another encoding the cognate trigger (Figure 4C). An additional set of CFPS powder samples were assembled containing a plasmid with constitutive LacZ expression as a positive control for LacZ reporter activity in PLGA. Lyophilized powder was mixed with acetone and PLGA solution and cast as films on glass in a multi-well slide. These films were rehydrated with a solution of Chlorophenol Red-β-D-Galactopyranoside (CPRG) indicator dye, which is cleaved by LacZ to change its color from yellow to magenta. CPRG color change activity was imaged over 8 h by an automated image scanner in a 37°C incubator. Constitutive production of LacZ was robust in PLGA, initiating the first recorded visible color change at 1 hour. The toehold switch-controlled reactions embedded in PLGA also performed as expected. Samples with only switch DNA had much slower color change compared to the samples with both switch and trigger DNA. The variability in switch performance compared to LacZ could be due to reduced overall expected LacZ productivity from the switch construct, combined with variable CFPS powder loading across multiple films. Future efforts to improve dispersion and uniformity in solvent-cast CFPS-PLGA could improve overall performance.

## 3. METHODOLOGY

### Reagents, DNA, strains

Unless otherwise noted, all reagents were purchased from Millipore Sigma, St. Louis, MO. The plasmid encoding sfGFP is pY71-sfGFP (GenBank accession MT346027).^71^ mRuby or LacZ coding sequences were inserted in place of sfGFP on the same plasmid backbone (pY71-mRuby, pY71-LacZ) for production of red fluorescence or LacZ enzyme, respectively.^52^ The toehold switch sensor was Switch H selected from a publication by Pardee et al^48^ and consisted of two plasmids pUCGA-swH-lacZ (switch) and pUCGA-trH (trigger) constructed as described by Blum et al.^52^ Plasmid DNA was purified from transformed *E. coli* using commercial midiprep or gigaprep kits from a variety of vendors.

Linear DNA encoding T7 polymerase-driven expression of colicin E1 with a 6xHis tag was amplified by a two-step PCR from plasmid pKSJ331 (a gift from Alan Finkelstein & Karen Jakes, Addgene plasmid #103063). The first PCR step used primers mslp151 and 153. The second PCR used primers mslp152 and 154 with the first PCR amplicon as template. Primer and final linear amplicon sequences are provided in Supplementary Table S2. The final amplicon solution was then treated with a purification kit (Nucleospin, Takara Bio Inc., San Jose, CA).

*E. coli* strains used in preparation of lysate were *E. coli* BL21 Star (DE3) for all experiments except for those where LacZ colorimetric reporter is expressed. Using the Rosetta 2(DE3) strain of *E. coli* (Novagen), a Δ*lacZα* derivative was constructed via CRISPR-Cas gene editing.^72^ A guide RNA sequence targeting *lacZ* (lacZb0344_68)^73^ and a homology-directed repair template corresponding to an in-frame deletion of base pairs 33–125 of the *lacZ* sequence and ∼500 base pairs of flanking DNA on each side were incorporated into plasmid pTargetF (a gift from Sheng Yang, Addgene plasmid #62226).^72^ The 93-base pair region of *lacZ* was deleted from the Rosetta 2(DE3) chromosome using this pTargetF derivative and plasmid pCas (a gift from Sheng Yang, Addgene plasmid #62225) using previously described methodology.^72^ The *lacZα* deletion and resulting loss of β-galactosidase activity were verified by DNA sequencing and plating on medium containing X-gal and IPTG.

### CFPS reaction preparation and lyophilization

*E. coli* lysates were prepared from flask cultures as described previously.^56^ T7 RNA polymerase was added exogenously after lysate preparation for all experiments. Lysis was performed either by sonication as described in detail previously^46^ or by homogenization using a Microfluidizer (Microfluidics M-110P). For homogenization the cell suspension (prepared the same as the sonication protocol) was processed in one pass followed by centrifugation and flash freezing at -80°C. A detailed protocol for microfluidizer operation is provided in the supplemental information.

CFPS reactions contained 30% v/v lysate and additional reagents following the PANOx-sp recipe as described in detail previously.^46^ All components and final concentrations are summarized in Supplementary Table S3. After mixing, CFPS reactions were lyophilized in plates, tubes, or vials depending on the scale described for specific experiments. CFPS reactions were mixed at normal aqueous concentrations at scales from 15 μL to 1 mL. “Small scale” volumes (< 100 μL) were deposited in polypropylene microplate wells or 1.5 mL microcentrifuge tubes. “Large scale” volumes (> 100 μL) were distributed in glass vials so that the height of the liquid did not exceed 2 cm.

The reactions were flash frozen in liquid nitrogen, then lyophilized using a shelf-type lyophilizer (SP Scientific, VirTis Wizard 2.0) with a 4 h primary drying step at shelf temperature -20°C, and an overnight (∼17 h) or longer (up to 48 h) secondary step with shelf temperature at 15°C. After removal from the lyophilizer, CFPS powders were stored for minimal time in a desiccator at room temperature if needed prior to treatment or rehydration. Untreated CFPS powder was rehydrated immediately after lyophilization and simultaneously with experiments to check activity for quality control. For CFPS material lyophilized in vials (> 100 μL starting volumes), the degree of water removal was measured by weight directly after lyophilization (Supplementary Table S1).

### Acetone and acetone-PLGA treatments in microplates

Small scale CFPS reaction solutions were distributed 15 μL per well into 96-well, v-bottom, polypropylene microplates (Costar 3357). The plates were lyophilized, then subjected to 15 μL acetone alone or a solution of 0.35 g/mL PLGA in acetone. The PLGA used was 75:25 L:G, 4-15k MW, acid terminated (Millipore Sigma 719919-5G). The plate was uncovered to allow acetone to evaporate for 1 h prior to rehydration with 15 μL water and measurement of protein production via fluorescence (sfGFP) or *E. coli* growth inhibition (colicin E1) via plate reader as described below.

### Acetone-cast PLGA films for DSLR camera imaging

Large 500 μL scale CFPS reactions were prepared with plasmids encoding sfGFP reporter (10 ng/μL) and lyophilized in vials. CFPS powder was combined with acetone-PLGA solution with a 3:1 volumetric ratio of PLGA solution to CFPS reaction solution pre-lyophilization. A disposable micropipette was used to coarsely crush and mix to form a suspension. 15 μL volumes of the suspension were drop-cast onto glass in an 8-well chamber slide (80821, Ibidi, Gräfelfing, Germany) and the acetone was allowed to evaporate for 1 hour at ambient conditions to form a CFPS-PLGA film.

Whole-film images were captured with a DSLR camera (Canon EOS Rebel T3, EF-S 18-55 mm IS II lens) mounted on a copy stand inside a darkened incubator set to 37°C. Canon EOS Rebel T3 Utility v.2 software was used to remotely control the camera settings and set up timelapse experiments. White light images were taken before rehydration, and at the start and end of the incubation. 180 μL water was added to fully submerge and rehydrate each film and the films were incubated at 37°C to observe the appearance of protein product. sfGFP fluorescence was imaged by placing the well slide on a DarkReader blue light transilluminator (Clare Chemical Research) and using an orange light filter placed over the camera lens. The blue light transilluminator is connected to the camera flash via an Arduino microcontroller. Fluorescence images were taken every minute for 3 h, then endpoint images were taken after 22 h. Shooting mode was manual, focus was manual, aperture was f/8.0, ISO was 3200, shutter speed was 0.6 seconds.

### Acetone-cast PLGA films for microscopy imaging

Small scale CFPS reactions were prepared with plasmids encoding the reporters sfGFP (10 ng/μL), mRuby (10 ng/μL), or water (no DNA control). Reactions were frozen in liquid nitrogen and lyophilized overnight in 1.5 mL microcentrifuge tubes. PLGA was dissolved in acetone (0.35 g/mL) and was added to lyophilized CFPS at a 1:1 volumetric ratio relative to the volume of pre-lyophilized CFPS reaction. A polypropylene pestle (F19923-0000, Bel-Art, South Wayne, NJ) was used to crush the CFPS powder into a uniform suspension in the presence of PLGA solution. 10 μL of each CFPS-PLGA mixture were cast into individual films in an 8-well chamber slide (80821, Ibidi, Gräfelfing, Germany). For dual function films, 10 μL of sfGFP CFPS-PLGA and 10 μL of mRuby CFPS-PLGA were cast side-by-side. The PLGA was allowed to cure uncovered for 4 h at room temperature. To detect fluorescence activity of CFPS embedded PLGA, 200 μL of nuclease free water was added to each well and slides were imaged by microscope over 24 h as described above. Fluorescence profiles for dual color films (Figure 3C) were measured in the Zen Blue software by drawing a rectangular area of interest in the profile tool encompassing the whole film, arrow pointing from left to right. Raw intensity values are plotted vs position in mm. For the graphed time course data (Figure 3E, F), Fluorescence intensity for entire film regions were measured in Zeiss Zen Blue software by drawing an area of interest to cover the entire film. sfGFP signal and mRuby signal intensities were integrated over the area at each time point for each sample. Graphed time course results for each condition show the averaged intensity value from 3 separate replicated films, with the time zero fluorescence values subtracted as background.

### PLGA-CFPS films with toehold switch and LacZ reporter

Lysate used with LacZ CFPS was prepared from Δ*lacZα* Rosetta 2(DE3) *E. coli*. 5 μg switch DNA, and either 10 μg trigger DNA (+Switch +Trigger) or equal volume PCR-grade water (+Switch -Trigger) were added to each 40 μL reaction volume. A positive control contained 500 ng plasmid that constitutively expressed LacZ (pY71-LacZ). A negative control did not contain DNA. 40uL of each thoroughly mixed reaction (CFPS+DNA) was aliquoted into individual 1.5 mL tubes, flash frozen in liquid nitrogen, and lyophilized with a shelf lyophilizer as described above (−20 ° 4 h primary, +15 ° overnight secondary). The following day, lyophilized samples were ground into a powder using a plastic pestle. Separately, PLGA (75:25) was completely dissolved in acetone (33% w/v) within a small glass bottle. 30μL of dissolved polymer was then added to each tube containing the crushed CFPS reactions and gently mixed by pipetting. 25 μL of each mixture was then pipetted into individual wells of an 8-well acetone-resistant plate (Ibidi). The PLGA was allowed to cure uncovered for 2 h at room temperature. Following evaporation, each reaction was rehydrated with 250 μL of 2 mM CPRG and incubated at 37 ° for 16 h. Images were taken every 30 minutes using an Epson Perfection V600 scanner placed inside of the incubator, with automated imaging controlled via custom software.^74^

### Heat-cast PCL films

A heat press (Dulytek DM800) was used to press PCL pellets (Mn 80,000, Millipore Sigma 440744) for 5 seconds at 80°C. PCL pellets weighed on average 14 mg ±1.9 mg. 30 μL aqueous CFPS reaction was placed on top of a PCL disc then frozen in a -80°C freezer. A shelf-type lyophilizer was used to dry the CFPS on PCL as described above. The second PCL disc was placed directly on top of the lyophilized material and re-pressed to embed the CFPS powder between melted PCL layers. Cut and whole PCL sandwich discs were placed in a 12-well microplate with coverslip-bottom wells (Cellvis P12-1.5H-N 12-well plate with #1.5HP glass bottom). The PCL pieces were covered with a 1% agarose solution in deionized water maintained at 50°C. The agarose was allowed to solidify for several minutes at 4°C. An agarose overlay was necessary to fully immobilize the PCL pieces for time course imaging since they did not adhere to the coverslip. Microscopy was performed over a 36 h time course as described below using mRuby and DIC imaging channels. The quantification of mRuby fluorescence signal measured by microscopy was performed using Zen Blue software (Carl Zeiss Microscopy, LLC). Quantification and normalization are further described in the supplemental information.

### Microscopy Imaging

Microscopy experiments used a Zeiss Axio observer Z1 inverted microscope with a Plan-Apochromat 10x/0.45 M27 objective, and incubation cabinet set to 37°C, with automated stage and focusing. Images were captured with an Axiocam 506 camera. Light source was Colibri 7 with a 475 nm LED used for sfGFP and a 567 nm LED for mRuby. Tiled images of each polymer sample were taken using up to three channels: DIC (106.6 ms exposure), sfGFP (ex: 480 nm, em: 505 nm, 360 ms exposure, 9% light intensity), and mRuby (ex: 577, em: 603, 5.5 s exposure, 100% light intensity). Images were taken each hour for the experiment durations described.

### Microplate reader fluorescence measurements

Immediately after rehydration, plates were sealed with a polypropylene mat (Costar 2080), transferred to a BioTek Synergy Neo2 microplate reader, and incubated at 37°C for 8 h. Formation of sfGFP fluorescence was monitored with ex/em: 485/528 nm. sfGFP readings in relative fluorescence units (RFU) were converted to equivalent concentration of fluorescein sodium salt (FSS) by measuring standard solutions: 0–25 μM of fluorescein sodium salt (CAS 518-47-8) in 100 mM sodium borate buffer solution, pH 9.5. mRuby fluorescence was monitored with ex/em: 558/605 nm and is reported in RFU. For all readings the extended gain function was selected.

### Colicin expression and growth inhibition assay

*E. coli* MG1655 cultures were prepared to assay colicin antimicrobial activity. On day 1 of the experiment, the strain was streaked from freezer stock onto LB agar. Then on day 2, colonies were used to inoculate 3 biological replicates in 5 mL LB media with 1% glucose in 50 mL conical tubes. These starter cultures were grown overnight at 37°C, and 250 rpm orbital shaking. On day 3, *E. coli* starter cultures were diluted 1:100 in LB + 1% glucose and grown until an OD600 of 0.05 or 4x108 cells/mL. Then the *E. coli* subcultures were diluted to ∼104 cells/mL and distributed at 200 μL per well in a sterile Costar 96-well flat bottom plate (#3370). All unused wells were filled with 200 μL blank media.

On day 2 of the experiment, linear colicin DNA was added at 40 nM final concentration to 15 μL CFPS reactions in a 96-well microplate prior to lyophilization. GamS (New England Biolabs, P0774S) was also added at a concentration of 0.3 μL per 15 μL reaction. When lyophilization was complete on day 3, dried reactions with and without DNA were exposed to acetone, a solution of PLGA in acetone, or left untreated. After solvent evaporation, reactions were rehydrated with 15 μL water and incubated for 2 h at 37°C to stimulate production of colicin. 10 μL aliquots of CFPS reactions were added to each well of the prepared microplate of diluted *E. coli* cultures.

The *E. coli* culture plate was sealed with a Breathe-Easy oxygen permeable plate sealer (Sigma Z380059). The plate was incubated at 30°C in a BioTek Synergy Neo2 microplate reader for 24 h with linear shaking at 567 cpm, a 1°C temperature gradient to prevent condensation and evaporation, and the optical density at 600 nm was read every 2 minutes.

## 4. DISCUSSION

Overall, this work demonstrates the potential of CFPS to bring DNA-encoded biological functionality to polymer materials. Lyophilized CFPS powder can tolerate both organic solvent and high temperature polymer processing conditions. Further, reactions embedded within polymers PLGA or PCL can be hydrated to carry out a variety of DNA-encoded functions. Formulating bioactivity into polymer composites has far-reaching connotations for both engineering new material functions and modulating the characteristics of the embedded biological systems.

Casting CFPS into polymer materials is a significant step beyond embedding individual enzymes, offering DNA-programmable functionality of a ubiquitous class of materials. This study confirms several CFPS functionalities remain active with the polymer-embedding technique, including constitutive protein synthesis of multiple protein types, use of linear or plasmid DNA, and sensor responsiveness. Future work could explore more complex functions such as logic gates; expression of proteins that degrade, repair, or otherwise manipulate the material; localized pattern generation; or combinations of multiple functions. The potential to implement any of these functions by simply changing the DNA offers significant flexibility. Compared to preserving the function of individual enzymes by painstakingly optimizing formulations or protein sequence each time, expressing a different protein of interest from a single optimized formulation for CFPS is dramatically simpler.

Differences in the hydrophobicity of PCL and PLGA polymers yielded different reactivation behaviors for CFPS. This highlights the potential for polymers to control hydration of the CFPS reaction in a material. One could envision using principles similar to controlled release to control hydration and therefore activate more and more of the CFPS reagents in the material over time. This might be used to extend the overall lifetime of CFPS-driven function in a polymer-CFPS system. Polymer barriers to water infiltration might also be used to extend shelf life. Thirdly, as the heat-cast PCL demonstrates, CFPS could be used to sense and/or repair damage to a material.

Aside from control over water transport, other small molecule transport could be slowed or blocked by the surrounding polymer matrix. Controlled transport could be used multiple ways. A material might allow the passage of an analyte of interest for sensing, while preventing infiltration of inhibitory contaminants. Substrates or intermediates might be concentrated locally to enhance the function of an enzyme cascade. Signaling between different bioparticles in a manner similar to quorum sensing could lead to long-range or patterned responses to stimuli within a material.

The sought-after concept of engineered living materials encompasses the use of living cells to endow anthropogenic materials with new features common in biology, including sensing, self-healing, and dynamic multi-functionality. Complex cell-free systems, though non-living, could be utilized to achieve many of these goals in polymer materials without the challenges associated with sustaining cell viability or concerns over release of genetically modified organisms. New form factors for CFPS can move applications toward commonplace materials and devices and away from highly technical lab uses. We anticipate that future development could include more classes of polymer materials with different properties, advanced formulations of dry cell-free material, advanced casting techniques like fiber spinning or layered deposition, and deeper understanding of the effects of hydration and polymer properties on biomolecule function and transport, all ultimately leading to useful real-world applications.

## ACKNOWLEDGMENTS

Funding was provided by the U.S. Army via the Chemical Biological Advanced Materials and Manufacturing Science Program (PE 0601102A Project VR9) at the Combat Capabilities Development Command (DEVCOM) Chemical Biological Center. Figure 3A was made in BioRender.com.

## Notes

### Competing Interest Statement

The authors have declared no competing interest.

