## Supplemental Information for "Cell-Free Protein Expression in Polymer Materials"

### Supplementary Information

#### *Microfluidizer Lysis Protocol*

- Turn Power on (left)
- Turn Motor on (front)
- Pack tank surrounding the cooling coil with ice/water – keep extra ice on hand as it melts.
- To process liquid in reservoir, pull the intensifier (black knob) to start, push to stop.
  - The psi should be between 15000–20000 (adjust using the process pressure adjustment knob).
  - When liquid disappears from the reservoir, that pressure pulse is referred to as pulse count 1, can run for at most two more pulse counts before intensifier should be turned off to avoid running air through the system.
- System is stored in ethanol or isopropyl alcohol. Pulse intensifier to run out ethanol/alcohol – collect outflow in container for disposal.
- Run DI water through until you see no alcohol – collect for disposal (2–2.5 reservoir runs).
- Run one more reservoir of water – collect for disposal.
- Run 50 mL of chosen buffer to prime system (for lysates use the same S30A buffer described in sonication protocol) and allow reservoir to empty – collect outflow for disposal.
- Prepare several clean 50 mL conical tubes for lysate collection, then add cell suspension to reservoir and start intensifier. It is recommended to collect the outflow in fractions with 2 pulses per fraction. Start keeping fractions collected from the time outflow first appears cloudy.
- When reservoir is empty of cell suspension, add 50 mL buffer to wash out (can stop intensifier to do this) – the first pulse or two after addition of wash buffer may still have material you can save.
- Stop collecting fractions when the lysate appears to become significantly more dilute.
- Run water though system (at least one full reservoir) – collect for waste.
- Take glass reservoir off machine and wash with soap and water.
- Run one tube of ethanol/alcohol to clean – collect for waste.

- Refill tube with ethanol/alcohol and run halfway to store system in alcohol.
- Turn off machine in reverse order (motor first then power source).
- Drain water from cooling coil tank and dry.

~20 mL of *E. coli* cell suspension typically yields four 2-pulse-fractions with good CFPS activity. Fractions are typically pooled before freezing.

Lysate is centrifuged, aliquoted, and flash frozen in the same manner as the sonicated lysate described previously.

***Notes on Preparation and Handling of Lyophilized CFPS Powder***

Batches of lyophilized CFPS reaction material up to 1 mL were prepared for solvent-casting experiments. At these larger scales, the performance of lyophilized batches was variable, and water absorbance during transfers was common when ambient humidity was high. Laboratory humidity control as well as quality checks like dry weight and activity controls are critical for a successful process. It was determined that dry weights near 80 mg per mL of starting reaction volume were typical for shelf-stable samples using this formulation. Dried samples were stored in a desiccator at room temperature. Samples that dried well and were stored properly had preserved activity for up to 10 weeks (Supplementary Table S1). Samples that absorbed water exhibited clumping, adhered to the vial and other surfaces, and changed color from white to yellow. After lyophilization, vials contain a dry cake that may be ground to form a powder. Initially, grinding was done open to air and powder was weighed into a vial containing polymer solution (Figure 1D). This only worked to any degree with very low ambient humidity, and still resulted in high variability between samples. Better reproducibility was possible by adding polymer solution directly to the lyophilized cake and grinding in the presence of acetone (Figure 3).

***Normalization Equation for CFPS-PCL Microscopy Measurements***

Regions of interest were drawn encompassing all of the film or film pieces, and the intensity sum for the mRuby channel was calculated for each region at each timepoint (grey values of the region are added together). The intensity sum values were normalized via the below equation and plotted in Figure 2C.

$$y = \frac{x_t - x_0}{Maximum\ x - x_0\ value}$$

$y$: normalized fluorescence value at timepoint t, arbitrary units

$x_t$: raw integrated sum value at time point t

$x_0$: raw integrated sum value at the initial time point. This is background fluorescence signal for the region of interest.

$Maximum\ x - x_0\ value$ : the background-subtracted integrated sum value for the region and timepoint with the highest fluorescent reading in the dataset (+ DNA cut film, 18 h timepoint for this dataset). This sets the highest value to 1 in the graph for easier comparison.

**Supplementary Figures**

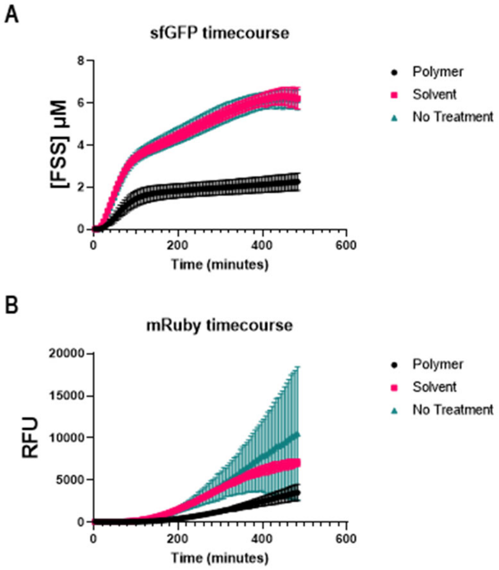


**Figure S1.** Time course data corresponding endpoint bar charts in Figure 1B and C of the main text. (A) sfGFP fluorescence signal in concentration of equivalent fluorescein [FSS] for samples containing pY71 sfGFP plasmid DNA. (B) mRuby fluorescence signal in RFU for samples containing pY71 mRuby plasmid DNA. Samples were treated with polymer (PLGA in acetone), solvent (acetone), or untreated. Error bars represent the 95% confidence interval, with $n \geq 6$.

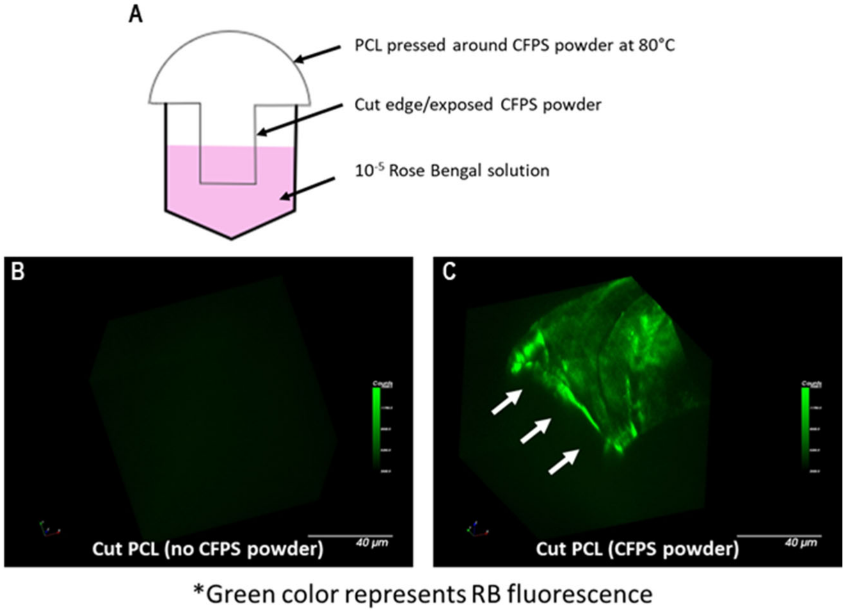


**Figure S2.** Infiltration of aqueous dye solution into cut PCL. (A) Diagram showing experiment setup with PCL cut to sit on top of a microplate well with cut edge partially submerged in an aqueous solution of Rose Bengal (RB) dye. (B) Microscopy showing that RB does not infiltrate cut PCL that does not contain any CFPS powder. (C) Microscopy showing RB has infiltrated the PCL containing CFPS powder. White arrows indicate the cut edge of the material. Scale bars are 40 μm.

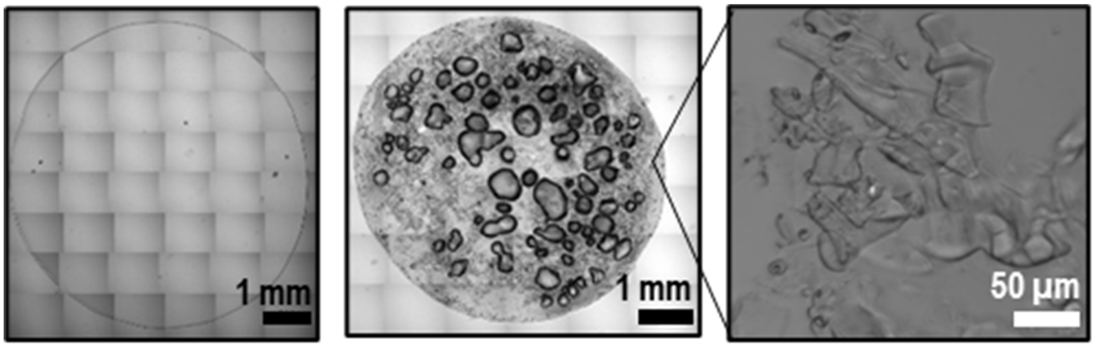


**Figure S3.** DIC microscopy. (left) Tiled DIC images depicting a PLGA film with no CFPS powder, (middle) Tiled DIC images depicting an entire CFPS-PLGA film, (right) Increased magnification DIC microscopy image to depict morphology of dried CFPS reaction powder embedded in PLGA solvent-cast film.

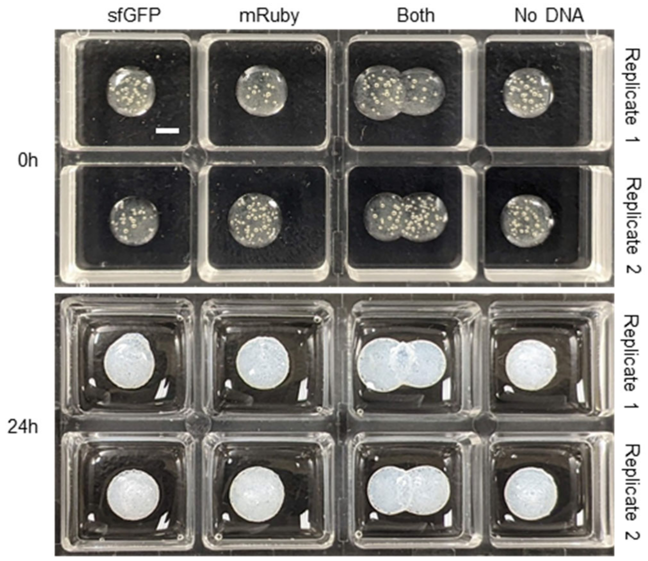


**Figure S4.** Additional replicates: PLGA films containing CFPS powder before and 24 h after rehydration. Bubbles can be seen in cast films following drying. Exposure to water causes PLGA to undergo hydrolysis, turning the films an opaque white color. Scale bar is 2 mm.

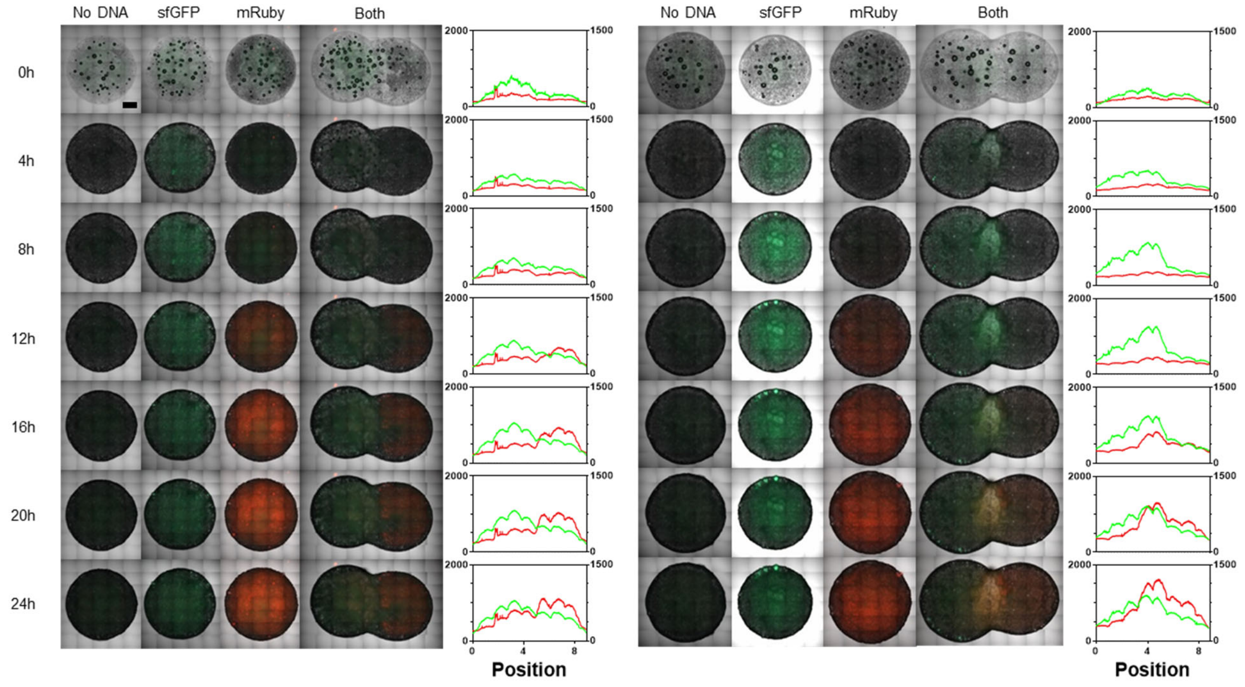


**Figure S5.** Additional replicates: CFPS of fluorescent reporters in PLGA over time. Films were prepared and imaged the same way as films in Figure 3C of the main text. Scale bar is 1 mm.

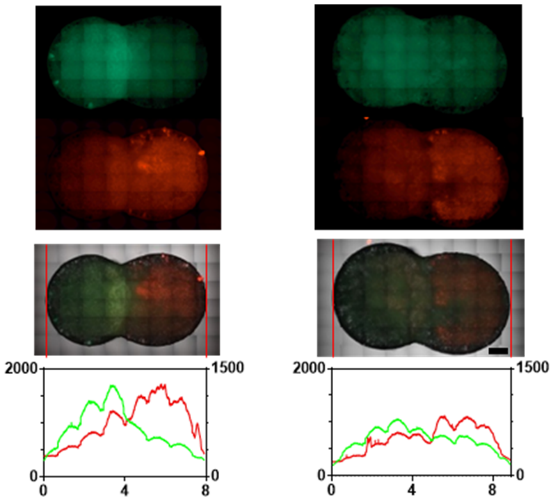


**Figure S6.** Two-color films endpoints: additional side-by-side comparison of the endpoints for three replicate two-color PLGA films with unmerged fluorescence channels. These are the same films depicted in Figure 3C of the main text and supplemental Figure S6. Each column of images are data for one replicate. Row labels from the top: sfGFP channel alone, mRuby channel alone, merged channels image (sfGFP, mRuby, and DIC), and profile data for sfGFP and mRuby fluorescence intensity. Red lines on merged microscopy images reflect the leftmost and rightmost positions measured in the profile graph.

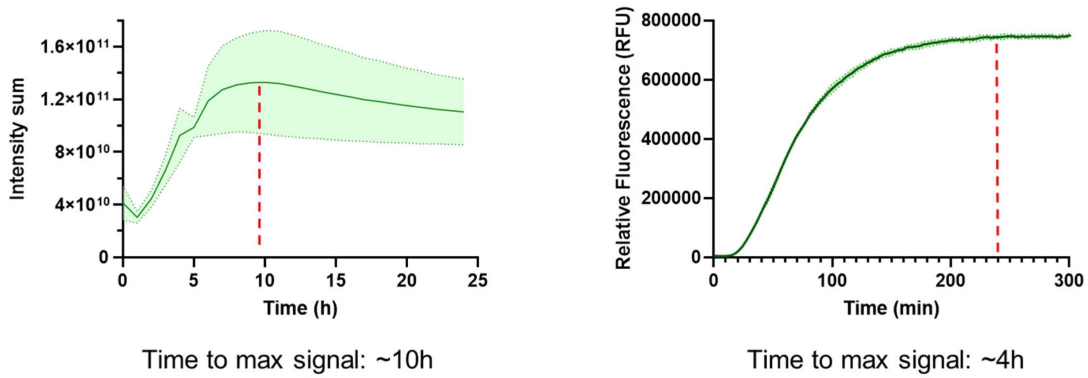


**Figure S7.** Dynamic measurements of sfGFP fluorescence in a film (left, measured by microscopy) or in solution (right, measured by microplate reader). Red dotted line indicates time to reach max fluorescence signal.

### Supplementary Tables

**Table S1.** Catalogue of shelf-life tests for lyophilized samples

| Sample | Date prepared | Protein product | Dry weight per mL (mg) | Age at shelf-life test (days) | Mass (mg) | Rehydration volume (µL) | Activity compared to Q.C. post-lyo (%) |
|---|---|---|---|---|---|---|---|
| 1 | 12/20/2021 | mRuby | 84.2 | 70 | 109.5 | 50 | 109.5 |
| 2 | 12/20/2021 | sfGFP | 83.6 | 70 | 101.1 | 40 | 101.1 |
| 3 | 12/27/2021 | mRuby | 67 | 63 | 119.4 | 40 | 119.4 |
| 4 | 12/27/2021 | mRuby | 67 | 63 | 133.7 | 40 | 133.7 |
| 5 | 12/27/2021 | mRuby | 67.2 | 63 | 89.5 | 50 | 89.5 |
| 6 | 12/27/2021 | mRuby | 73.6 | 63 | 82.9 | 50 | 82.9 |
| 7 | 12/27/2021 | sfGFP | 67.5 | 63 | 102.9 | 30 | 102.9 |
| 8 | 1/12/2022 | sfGFP | 86.56 | 47 | 20.7 | 140 | 20.7 |
| 9 | 1/12/2022 | sfGFP | 87.05 | 47 | 41.4 | 50 | 41.4 |
| 10 | 1/12/2022 | mRuby | 87.12 | 47 | 177.1 | 60 | 177.1 |
| 11 | 1/21/2022 | mRuby | 86.72 | 47 | 112.9 | 50 | 112.9 |

**Table S2.** DNA constructs from this study

| Name, description | Sequence |
|---|---|
| pY71 GFP – plasmid encoding T7 promoter expression of sfGFP, Kanamycin resistance, ColE1 origin | GGATCCTGCAGTTGAGATCCTTTTTTTCTGCGCGTAATCTGCTGCTTGCAAACAAAAAAAC<br>CACCGCTACCAGCGGTGGTTTGTTTGCCGGATCAAGAGCTACCAACTCTTTTTCCGAAGGT<br>AACTGGCTTCAGCAGAGCGCAGATACCAAATACTGTCCTTCTAGTGTAGCCGTAGTTAGG<br>CCACCACTTCAAGAACTCTGTAGCACCGCCTACATACCTCGCTCTGCTAATCCTGTTACCA<br>GTGGCTGCTGCCAGTGGCGATAAGTCGTGTCTTACCGGGTTGGACTCAAGACGATAGTTA<br>CCGGATAAGGCGCAGCGGTCGGGCTGAACGGGGGGTTCGTGCACACAGCCCAGCTTGGA<br>GCGAACGACCTACACCGAACTGAGATACCTACAGCGTGAGCATTGAGAAAGCGCCACGC<br>TTCCCGAAGGGAGAAAGGCGGACAGGTATCCGGTAAGCGGCAGGGTCGGAACAGGAGAG<br>CGCACGAGGGAGCTTCCAGGGGGAAACGCCTGGTATCTTTATAGTCCTGTCGGGTTTCGC<br>CACCTCTGACTTGAGCGTCGATTTTTGTGATGCTCGTCAGGGGGGCGGAGCCTATGGAAA<br>CGAATTCAGATCTCGATCCCGCGAAATTAATACGACTCACTATAGGGAGACCACAACGGT<br>TTCCCTCTAGAAATAATTTTGTTTAACTTTAAGAAGGAGATATACATATGAGCAAAGGTG<br>AAGAACTGTTTACCGGCGTTGTGCCGATTCTGGTGGAACTGGATGGCGATGTGAACGGTC<br>ACAAATTCAGCGTGCGTGGTGAAGGTGAAGGCGATGCCACGATTGGCAAACTGACGCTG<br>AAATTTATCTGCACCACCGGCAAACTGCCGGTGCCGTGGCCGACGCTGGTGACCACCCTG<br>ACCTATGGCGTTCAGTGTTTTAGTCGCTATCCGGATCACATGAAACGTCACGATTTCTTTA<br>AATCTGCAATGCCGGAAGGCTATGTGCAGGAACGTACGATTAGCTTTAAAGATGATGGCA<br>AATATAAAACGCGCGCCGTTGTGAAATTTGAAGGCGATACCCTGGTGAACCGCATTGAAC<br>TGAAAGGCACGGATTTTAAAGAAGATGGCAATATCCTGGGCCATAAACTGGAATACAACT<br>TTAATAGCCATAATGTTTATATTACGGCGGATAAACAGAAAAATGGCATCAAAGCGAATT<br>TTACCGTTCGCCATAACGTTGAAGATGGCAGTGTGCAGCTGGCAGATCATTATCAGCAGA<br>ATACCCCGATTGGTGATGGTCCGGTGCTGCTGCCGGATAATCATTATCTGAGCACGCAGA<br>CCGTTCTGTCTAAAGATCCGAACGAAAAACGGGACCACATGGTTCTGCACGAATATGTGA<br>ATGCGGCAGGTATTACGTGGAGCCATCCGCAGTTCGAAAAATAATAAGTCGACCGGCTGC<br>TAACAAAGCCCGAAAGGAAGCTGAGTTGGCTGCTGCCACCGCTGAGCAATAACTAGCATA<br>ACCCCTTGGGGCCTCTAAACGGGTCTTGAGGGGTTTTTTGCTGAAAGCCAATTCTGATTAG<br>AAAAACTCATCGAGCATCAAATGAAACTGCAATTTATTCATATCAGGATTATCAATACCA<br>TATTTTTGAAAAAGCCGTTTCTGTAATGAAGGAGAAAACTCACCGAGGCAGTTCCATAGG<br>ATGGCAAGATCCTGGTATCGGTCTGCGATTCCGACTCGTCCAACATCAATACAACCTATTA<br>ATTTCCCCTCGTCAAAAATAAGGTTATCAAGTGAGAAATCACCATGAGTGACGACTGAAT<br>CCGGTGAGAATGGCAAAAGCTTATGCATTTCTTTCCAGACTTGTTCAACAGGCCAGCCATT<br>ACGCTCGTCATCAAAATCACTCGCATCAACCAAACCGTTATTCATTCGTGATTGCGCCTGA<br>GCGAGACGAAATACGCGATCGCTGTTAAAAGGACAATTACAAACAGGAATCGAATGCAA<br>CCGGCGCAGGAACACTGCCAGCGCATCAACAATATTTTCACCTGAATCAGGATATTCTTCT<br>AATACCTGGAATGCTGTTTTCCCGGGGATCGCAGTGGTGAGTAACCATGCATCATCAGGA |

| | |
|---|---|
| | GTACGGATAAAATGCTTGATGGTCGGAAGAGGCATAAATTCCGTCAGCCAGTTTAGTCTG<br>ACCATCTCATCTGTAACATCATTGGCAACGCTACCTTTGCCATGTTTCAGAAACAACTCTG<br>GCGCATCGGGCTTCCCATACAATCGATAGATTGTCGCACCTGATTGCCCGACATTATCGCG<br>AGCCCATTTATACCCATATAAATCAGCATCCATGTTGGAATTTAATCGCGGCTTCGAGCAA<br>GACGTTTCCCGTTGAATATGGCTCATAACACCCCTTGTATTACTGTTTATGTAAGCAGACA<br>GTTTTATTGTTCATGATGATATATTTTTATCTTGTGCAATGTAACATCAGAGATTTTGAGAC<br>ACAACGT |
| pY71 mRuby2 – plasmid encoding T7 expression of mRuby, Kanamycin resistance, ColE1 origin | GGATCCTGCAGTTGAGATCCTTTTTTTCTGCGCGTAATCTGCTGCTTGCAAACAAAAAAAC<br>CACCGCTACCAGCGGTGGTTTGTTTGCCGGATCAAGAGCTACCAACTCTTTTTCCGAAGGT<br>AACTGGCTTCAGCAGAGCGCAGATACCAAATACTGTCCTTCTAGTGTAGCCGTAGTTAGG<br>CCACCACTTCAAGAACTCTGTAGCACCGCCTACATACCTCGCTCTGCTAATCCTGTTACCA<br>GTGGCTGCTGCCAGTGGCGATAAGTCGTGTCTTACCGGGTTGGACTCAAGACGATAGTTA<br>CCGGATAAGGCGCAGCGGTCGGGCTGAACGGGGGGTTCGTGCACACAGCCCAGCTTGGA<br>GCGAACGACCTACACCGAACTGAGATACCTACAGCGTGAGCATTGAGAAAGCGCCACGC<br>TTCCCGAAGGGAGAAAGGCGGACAGGTATCCGGTAAGCGGCAGGGTCGGAACAGGAGAG<br>CGCACGAGGGAGCTTCCAGGGGGAAACGCCTGGTATCTTTATAGTCCTGTCGGGTTTCGC<br>CACCTCTGACTTGAGCGTCGATTTTTGTGATGCTCGTCAGGGGGGCGGAGCCTATGGAAA<br>CGAATTCAGATCTCGATCCCGCGAAATTAATACGACTCACTATAGGGAGACCACAACGGT<br>TTCCCTCTAGAAATAATTTTGTTTAACTTTAAGAAGGAGATATACATATGGTGTCCAAAGG<br>AGAGGAGTTAATCAAGGAAAACATGAGAATGAAAGTTGTCATGGAGGGCTCCGTTAATG<br>GTCACCAATTCAAGTGTACAGGGGAAGGTGAAGGTAATCCTTACATGGGTACACAAACTA<br>TGAGAATTAAAGTAATTGAAGGCGGACCACTACCATTTGCATTTGACATTCTGGCAACGT<br>CATTCATGTACGGATCACGAACTTTCATCAAGTACCCTAAAGGTATACCAGACTTTTTCAA<br>GCAATCTTTTCCAGAGGGTTTTACATGGGAAAGGGTTACAAGATACGAAGATGGGGGTGT<br>CGTCACAGTTATGCAAGATACTTCATTAGAAGATGGCTGCCTTGTCTATCATGTGCAAGTA<br>AGAGGGGTGAATTTTCCTTCTAACGGACCTGTGATGCAGAAAAAGACCAAAGGTTGGGAA<br>CCAAATACTGAAATGATGTACCCAGCTGATGGAGGTTTGAGAGGCTACACACACATGGCG<br>CTTAAAGTTGATGGTGGAGGTCATTTGTCTTGTAGTTTTGTTACCACTTATCGTTCTAAAA<br>AGACTGTTGGCAATATCAAAATGCCAGGAATACATGCTGTAGACCACAGACTAGAAAGA<br>CTCGAAGAGAGCGATAACGAAATGTTCGTTGTACAGAGAGAGCATGCCGTAGCCAAATTT<br>GCTGGCTTAGGCGGTGGTATGGATGAATTGTATAAGTGATAAGTCGACCGGCTGCTAACA<br>AAGCCCGAAAGGAAGCTGAGTTGGCTGCTGCCACCGCTGAGCAATAACTAGCATAACCCC<br>TTGGGGCCTCTAAACGGGTCTTGAGGGGTTTTTTGCTGAAAGCCAATTCTGATTAGAAAA<br>ACTCATCGAGCATCAAATGAAACTGCAATTTATTCATATCAGGATTATCAATACCATATTT<br>TTGAAAAAGCCGTTTCTGTAATGAAGGAGAAAACTCACCGAGGCAGTTCCATAGGATGGC<br>AAGATCCTGGTATCGGTCTGCGATTCCGACTCGTCCAACATCAATACAACCTATTAATTTC<br>CCCTCGTCAAAAATAAGGTTATCAAGTGAGAAATCACCATGAGTGACGACTGAATCCGGT<br>GAGAATGGCAAAAGCTTATGCATTTCTTTCCAGACTTGTTCAACAGGCCAGCCATTACGCT<br>CGTCATCAAAATCACTCGCATCAACCAAACCGTTATTCATTCGTGATTGCGCCTGAGCGAG<br>ACGAAATACGCGATCGCTGTTAAAAGGACAATTACAAACAGGAATCGAATGCAACCGGC<br>GCAGGAACACTGCCAGCGCATCAACAATATTTTCACCTGAATCAGGATATTCTTCTAATAC<br>CTGGAATGCTGTTTTCCCGGGGATCGCAGTGGTGAGTAACCATGCATCATCAGGAGTACG<br>GATAAAATGCTTGATGGTCGGAAGAGGCATAAATTCCGTCAGCCAGTTTAGTCTGACCAT<br>CTCATCTGTAACATCATTGGCAACGCTACCTTTGCCATGTTTCAGAAACAACTCTGGCGCA<br>TCGGGCTTCCCATACAATCGATAGATTGTCGCACCTGATTGCCCGACATTATCGCGAGCCC<br>ATTTATACCCATATAAATCAGCATCCATGTTGGAATTTAATCGCGGCTTCGAGCAAGACGT<br>TTCCCGTTGAATATGGCTCATAACACCCCTTGTATTACTGTTTATGTAAGCAGACAGTTTT<br>ATTGTTCATGATGATATATTTTTATCTTGTGCAATGTAACATCAGAGATTTTGAGACACAA<br>CGT |
| pY71 LacZ – plasmid encoding T7 expression of LacZ, Kanamycin resistance, ColE1 origin | TAATAAACAATAACTGAATAGGGATCCCGACTGGCGAGAGCCAGGTAACGAATGGATCC<br>TTCAGCCAAAAAACTTAAGACCGCCGGTCTTGTCCACTACCTTGCAGTAATGCGGTGGAC<br>AGGATCGGCGGTTTTCTTTTCTCTTCTCAAGCTGAAAGCCAATTCTGATTAGAAAAACTCA<br>TCGAGCATCAAATGAAACTGCAATTTATTCATATCAGGATTATCAATACCATATTTTTGAA<br>AAAGCCGTTTCTGTAATGAAGGAGAAAACTCACCGAGGCAGTTCCATAGGATGGCAAGAT<br>CCTGGTATCGGTCTGCGATTCCGACTCGTCCAACATCAATACAACCTATTAATTTCCCCTC<br>GTCAAAAATAAGGTTATCAAGTGAGAAATCACCATGAGTGACGACTGAATCCGGTGAGA<br>ATGGCAAAAGCTTATGCATTTCTTTCCAGACTTGTTCAACAGGCCAGCCATTACGCTCGTC<br>ATCAAAATCACTCGCATCAACCAAACCGTTATTCATTCGTGATTGCGCCTGAGCGAGACG<br>AAATACGCGATCGCTGTTAAAAGGACAATTACAAACAGGAATCGAATGCAACCGGCGCA<br>GGAACACTGCCAGCGCATCAACAATATTTTCACCTGAATCAGGATATTCTTCTAATACCTG<br>GAATGCTGTTTTCCCGGGGATCGCAGTGGTGAGTAACCATGCATCATCAGGAGTACGGAT<br>AAAATGCTTGATGGTCGGAAGAGGCATAAATTCCGTCAGCCAGTTTAGTCTGACCATCTC<br>ATCTGTAACATCATTGGCAACGCTACCTTTGCCATGTTTCAGAAACAACTCTGGCGCATCG<br>GGCTTCCCATACAATCGATAGATTGTCGCACCTGATTGCCCGACATTATCGCGAGCCCATT<br>TATACCCATATAAATCAGCATCCATGTTGGAATTTAATCGCGGCTTCGAGCAAGACGTTTC<br>CCGTTGAATATGGCTCATAACACCCCTTGTATTACTGTTTATGTAAGCAGACAGTTTTATT<br>GTTCATGATGATATATTTTTATCTTGTGCAATGTAACATCAGAGATTTTGAGACACAACGT<br>GGATCCTGCAGTTGAGATCCTTTTTTTCTGCGCGTAATCTGCTGCTTGCAAACAAAAAAAC<br>CACCGCTACCAGCGGTGGTTTGTTTGCCGGATCAAGAGCTACCAACTCTTTTTCCGAAGGT<br>AACTGGCTTCAGCAGAGCGCAGATACCAAATACTGTCCTTCTAGTGTAGCCGTAGTTAGG |

| | |
|---|---|
| | CCACCACTTCAAGAACTCTGTAGCACCGCCTACATACCTCGCTCTGCTAATCCTGTTACCA<br>GTGGCTGCTGCCAGTGGCGATAAGTCGTGTCTTACCGGGTTGGACTCAAGACGATAGTTA<br>CCGGATAAGGCGCAGCGGTCGGGCTGAACGGGGGGTTCGTGCACACAGCCCAGCTTGGA<br>GCGAACGACCTACACCGAACTGAGATACCTACAGCGTGAGCATTGAGAAAGCGCCACGC<br>TTCCCGAAGGGAGAAAGGCGGACAGGTATCCGGTAAGCGGCAGGGTCGGAACAGGAGAG<br>CGCACGAGGGAGCTTCCAGGGGGAAACGCCTGGTATCTTTATAGTCCTGTCGGGTTTCGC<br>CACCTCTGACTTGAGCGTCGATTTTTGTGATGCTCGTCAGGGGGGCGGAGCCTATGGAAA<br>CGAATTCAGATCTCGATCCCGCGAAATTAATACGACTCACTATAGGGAGCTGTCACCGGA<br>TGTGCTTTCCGGTCTGATGAGTCCGTGAGGACGAAACAGCCTCTACAAATAATTTTGTTTA<br>ATCTAGAGAAAGAGGAGAAATACTAGATGACCATGATTACGGATTCACTGGCCGTCGTTT<br>TACAACGTCGTGACTGGGAAAACCCTGGCGTTACCCAACTTAATCGCCTTGCAGCACATC<br>CCCCTTTCGCCAGCTGGCGTAATAGCGAAGAGGCCCGCACCGATCGCCCTTCCCAACAGT<br>TGCGCAGCCTGAATGGCGAATGGCGCTTTGCCTGGTTTCCGGCACCAGAAGCGGTGCCGG<br>AAAGCTGGCTGGAGTGCGATCTTCCTGAGGCCGATACTGTCGTCGTCCCCTCAAACTGGC<br>AGATGCACGGTTACGATGCGCCCATCTACACCAACGTGACCTATCCCATTACGGTCAATC<br>CGCCGTTTGTTCCCACGGAGAATCCGACGGGTTGTTACTCGCTCACATTTAATGTTGATGA<br>AAGCTGGCTACAGGAAGGCCAGACGCGAATTATTTTTGATGGCGTTAACTCGGCGTTTCA<br>TCTGTGGTGCAACGGGCGCTGGGTCGGTTACGGCCAGGACAGTCGTTTGCCGTCTGAATTT<br>GACCTGAGCGCATTTTTACGCGCCGGAGAAAACCGCCTCGCGGTGATGGTGCTGCGCTGG<br>AGTGACGGCAGTTATCTGGAAGATCAGGATATGTGGCGGATGAGCGGCATTTTCCGTGAC<br>GTCTCGTTGCTGCATAAACCGACTACACAAATCAGCGATTTCCATGTTGCCACTCGCTTTA<br>ATGATGATTTCAGCCGCGCTGTACTGGAGGCTGAAGTTCAGATGTGCGGCGAGTTGCGTG<br>ACTACCTACGGGTAACAGTTTCTTTATGGCAGGGTGAAACGCAGGTCGCCAGCGGCACCG<br>CGCCTTTCGGCGGTGAAATTATCGATGAGCGTGGTGGTTATGCCGATCGCGTCACACTAC<br>GTCTGAACGTCGAAAACCCGAAACTGTGGAGCGCCGAAATCCCGAATCTCTATCGTGCGG<br>TGGTTGAACTGCACACCGCCGACGGCACGCTGATTGAAGCAGAAGCCTGCGATGTCGGTT<br>TCCGCGAGGTGCGGATTGAAAATGGTCTGCTGCTGCTGAACGGCAAGCCGTTGCTGATTC<br>GAGGCGTTAACCGTCACGAGCATCATCCTCTGCATGGTCAGGTCATGGATGAGCAGACGA<br>TGGTGCAGGATATCCTGCTGATGAAGCAGAACAACTTTAACGCCGTGCGCTGTTCGCATT<br>ATCCGAACCATCCGCTGTGGTACACGCTGTGCGACCGCTACGGCCTGTATGTGGTGGATG<br>AAGCCAATATTGAAACCCACGGCATGGTGCCAATGAATCGTCTGACCGATGATCCGCGCT<br>GGCTACCGGCGATGAGCGAACGCGTAACGCGAATGGTGCAGCGCGATCGTAATCACCCG<br>AGTGTGATCATCTGGTCGCTGGGGAATGAATCAGGCCACGGCGCTAATCACGACGCGCTG<br>TATCGCTGGATCAAATCTGTCGATCCTTCCCGCCCGGTGCAGTATGAAGGCGGCGGAGCC<br>GACACCACGGCCACCGATATTATTTGCCCGATGTACGCGCGCGTGGATGAAGACCAGCCC<br>TTCCCGGCTGTGCCGAAATGGTCCATCAAAAAATGGCTTTCGCTACCTGGAGAGACGCGC<br>CCGCTGATCCTTTGCGAATACGCCCACGCGATGGGTAACAGTCTTGGCGGTTTCGCTAAAT<br>ACTGGCAGGCGTTTCGTCAGTATCCCCGTTTACAGGGCGGCTTCGTCTGGGACTGGGTGG<br>ATCAGTCGCTGATTAAATATGATGAAAACGGCAACCCGTGGTCGGCTTACGGCGGTGATT<br>TTGGCGATACGCCGAACGATCGCCAGTTCTGTATGAACGGTCTGGTCTTTGCCGACCGCAC<br>GCCGCATCCAGCGCTGACGGAAGCAAAACACCAGCAGCAGTTTTTCCAGTTCCGTTTATC<br>CGGGCAAACCATCGAAGTGACCAGCGAATACCTGTTCCGTCATAGCGATAACGAGCTCCT<br>GCACTGGATGGTGGCGCTGGATGGTAAGCCGCTGGCAAGCGGTGAAGTGCCTCTGGATGT<br>CGCTCCACAAGGTAAACAGTTGATTGAACTGCCTGAACTACCGCAGCCGGAGAGCGCCGG<br>GCAACTCTGGCTCACAGTACGCGTAGTGCAACCGAACGCGACCGCATGGTCAGAAGCCGG<br>GCACATCAGCGCCTGGCAGCAGTGGCGTCTGGCGGAAAACCTCAGTGTGACGCTCCCCGC<br>CGCGTCCCACGCCATCCCGCATCTGACCACCAGCGAAATGGATTTTTGCATCGAGCTGGG<br>TAATAAGCGTTGGCAATTTAACCGCCAGTCAGGCTTTCTTTCACAGATGTGGATTGGCGAT<br>AAAAAACAACTGCTGACGCCGCTGCGCGATCAGTTCACCCGTGCACCGCTGGATAACGAC<br>ATTGGCGTAAGTGAAGCGACCCGCATTGACCCTAACGCCTGGGTCGAACGCTGGAAGGCG<br>GCGGGCCATTACCAGGCCGAAGCAGCGTTGTTGCAGTGCACGGCAGATACACTTGCTGAT<br>GCGGTGCTGATTACGACCGCTCACGCGTGGCAGCATCAGGGGAAAACCTTATTTATCAGC<br>CGGAAAACCTACCGGATTGATGGTAGTGGTCAAATGGCGATTACCGTTGATGTTGAAGTG<br>GCGAGCGATACACCGCATCCGGCGCGGATTGGCCTGAACTGCCAGCTGGCGCAGGTAGCA<br>GAGCGGGTAAACTGGCTCGGATTAGGGCCGCAAGAAAACTATCCCGACCGCCTTACTGCC<br>GCCTGTTTTGACCGCTGGGATCTGCCATTGTCAGACATGTATACCCCGTACGTCTTCCCGA<br>GCGAAAACGGTCTGCGCTGCGGGACGCGCGAATTGAATTATGGCCCACACCAGTGGCGCG<br>GCGACTTCCAGTTCAACATCAGCCGCTACAGTCAACAGCAACTGATGGAAACCAGCCATC<br>GCCATCTGCTGCACGCGGAAGAAGGCACATGGCTGAATATCGACGGTTTCCATATGGGGA<br>TTGGTGGCGACGACTCCTGGAGCCCGTCAGTATCGGCGGAATTCCAGCTGAGCGCCGGTC<br>GCTACCATTACCAGTTGGTCTGGTGTCAAAAA |
| pUCGA-swH-lacZ – plasmid encoding T7 expression of LacZ translationally regulated by swH toehold switch | GGAAGTTTGTCTAGATCTCAGGCGTGGATGCGGGTACCGAGCTCGAATTCACTGGCCGTC<br>GTTTTACAACGTCGTGACTGGGAAAACCCTGGCGTTACCCAACTTAATCGCCTTGCAGCAC<br>ATCCCCCTTTCGCCAGCTGGCGTAATAGCGAAGAGGCCCGCACCGATCGCCCTTCCCAAC<br>AGTTGCGCAGCCTGAATGGCGAATGGCGCCTGATGCGGTATTTTCTCCTTACGCATCTGTG<br>CGGTATTTCACACCGCATATGGTGCACTCTCAGTACAATCTGCTCTGATGCCGCATAGTTA<br>AGCCAGCCCCGACACCCGCCAACACCCGCTGACGCGCCCTGACGGGCTTGTCTGCTCCCG<br>GCATCCGCTTACAGACAAGCTGTGACCGTCTCCGGGAGCTGCATGTGTCAGAGGTTTTCA<br>CCGTCATCACCGAAACGCGCGAGACGAAAGGGCCTCGTGATACGCCTATTTTTATAGGTT<br>AATGTCATGATAATAATGGTTTCTTAGACGTCAGGTGGCACTTTTCGGGGAAATGTGCGC |

| | |
|---|---|
| | GGAACCCCTATTTGTTTATTTTTCTAAATACATTCAAATATGTATCCGCTCATGAGACAAT<br>AACCCTGATAAATGCTTCAATAATATTGAAAAAGGAAGAGTATGAGTATTCAACATTTCC<br>GTGTCGCCCTTATTCCCTTTTTTGCGGCATTTTGCCTTCCTGTTTTTGCTCACCCAGAAACG<br>CTGGTGAAAGTAAAAGATGCTGAAGATCAGTTGGGTGCACGAGTGGGTTACATCGAACTG<br>GATCTCAACAGCGGTAAGATCCTTGAGAGTTTTCGCCCCGAAGAACGTTTTCCAATGATG<br>AGCACTTTTAAAGTTCTGCTATGTGGCGCGGTATTATCCCGTATTGACGCCGGGCAAGAGC<br>AACTCGGTCGCCGCATACACTATTCTCAGAATGACTTGGTTGAGTACTCACCAGTCACAG<br>AAAAGCATCTTACGGATGGCATGACAGTAAGAGAATTATGCAGTGCTGCCATAACCATGA<br>GTGATAACACTGCGGCCAACTTACTTCTGACAACGATCGGAGGACCGAAGGAGCTAACCG<br>CTTTTTTGCACAACATGGGGGATCATGTAACTCGCCTTGATCGTTGGGAACCGGAGCTGA<br>ATGAAGCCATACCAAACGACGAGCGTGACACCACGATGCCTGTAGCAATGGCAACAACG<br>TTGCGCAAACTATTAACTGGCGAACTACTTACTCTAGCTTCCCGGCAACAATTAATAGACT<br>GGATGGAGGCGGATAAAGTTGCAGGACCACTTCTGCGCTCGGCCCTTCCGGCTGGCTGGT<br>TTATTGCTGATAAATCTGGAGCCGGTGAGCGTGGGTCTCGCGGTATCATTGCAGCACTGG<br>GGCCAGATGGTAAGCCCTCCCGTATCGTAGTTATCTACACGACGGGGAGTCAGGCAACTA<br>TGGATGAACGAAATAGACAGATCGCTGAGATAGGTGCCTCACTGATTAAGCATTGGTAAC<br>TGTCAGACCAAGTTTACTCATATATACTTTAGATTGATTTAAAACTTCATTTTTAATTTAAA<br>AGGATCTAGGTGAAGATCCTTTTTGATAATCTCATGACCAAAATCCCTTAACGTGAGTTTT<br>CGTTCCACTGAGCGTCAGACCCCGTAGAAAAGATCAAAGGATCTTCTTGAGATCCTTTTTT<br>TCTGCGCGTAATCTGCTGCTTGCAAACAAAAAAACCACCGCTACCAGCGGTGGTTTGTTTG<br>CCGGATCAAGAGCTACCAACTCTTTTTCCGAAGGTAACTGGCTTCAGCAGAGCGCAGATA<br>CCAAATACTGTTCTTCTAGTGTAGCCGTAGTTAGGCCACCACTTCAAGAACTCTGTAGCAC<br>CGCCTACATACCTCGCTCTGCTAATCCTGTTACCAGTGGCTGCTGCCAGTGGCGATAAGTC<br>GTGTCTTACCGGGTTGGACTCAAGACGATAGTTACCGGATAAGGCGCAGCGGTCGGGCTG<br>AACGGGGGGTTCGTGCACACAGCCCAGCTTGGAGCGAACGACCTACACCGAACTGAGAT<br>ACCTACAGCGTGAGCTATGAGAAAGCGCCACGCTTCCCGAAGGGAGAAAGGCGGACAGG<br>TATCCGGTAAGCGGCAGGGTCGGAACAGGAGAGCGCACGAGGGAGCTTCCAGGGGGAAA<br>CGCCTGGTATCTTTATAGTCCTGTCGGGTTTCGCCACCTCTGACTTGAGCGTCGATTTTTGT<br>GATGCTCGTCAGGGGGGCGGAGCCTATGGAAAAACGCCAGCAACGCGGCCTTTTTACGGT<br>TCCTGGCCTTTTGCTGGCCTTTTGCTCACATGTTCTTTCCTGCGTTATCCCCTGATTCTGTG<br>GATAACCGTATTACCGCCTTTGAGTGAGCTGATACCGCTCGCCGCAGCCGAACGACCGAG<br>CGCAGCGAGTCAGTGAGCGAGGAAGCGGAAGAGCGCCCAATACGCAAACCGCCTCTCCC<br>CGCGCGTTGGCCGATTCATTAATGCAGCTGGCACGACAGGTTTCCCGACTGGAAAGCGGG<br>CAGTGAGCGCAACGCAATTAATGTGAGTTAGCTCACTCATTAGGCACCCCAGGCTTTACA<br>CTTTATGCTTCCGGCTCGTATGTTGTGTGGAATTGTGAGCGGATAACAATTTCACACAGGA<br>AACAGCTATGACCATGATTACGCCAAGCTTGCATGCCTGCAGGTCGACTCTAGAGGATCG<br>AGCACATCTCGTTCGCTATTCAGGGATTGTAATACGACTCACTATAGGGTGAATGAATTGT<br>AGGCTTGTTATAGTTATGAACAGAGGAGACATAACATGAACAAGCCTAACCTGGCGGCAG<br>CGCAAAAGATGACCATGATTACGGATTCACTGGCCGTCGTTTTACAACGTCGTGACTGGG<br>AAAACCCTGGCGTTACCCAACTTAATCGCCTTGCAGCACATCCCCCTTTCGCCAGCTGGCG<br>TAATAGCGAAGAGGCCCGCACCGATCGCCCTTCCCAACAGTTGCGCAGCCTGAATGGCGA<br>ATGGCGCTTTGCCTGGTTTCCGGCACCAGAAGCGGTGCCGGAAAGCTGGCTGGAGTGCGA<br>TCTTCCTGAGGCCGATACTGTCGTCGTCCCCTCAAACTGGCAGATGCACGGTTACGATGCG<br>CCCATCTACACCAACGTGACCTATCCCATTACGGTCAATCCGCCGTTTGTTCCCACGGAGA<br>ATCCGACGGGTTGTTACTCGCTCACATTTAATGTTGATGAAAGCTGGCTACAGGAAGGCC<br>AGACGCGAATTATTTTTGATGGCGTTAACTCGGCGTTTCATCTGTGGTGCAACGGGCGCTG<br>GGTCGGTTACGGCCAGGACAGTCGTTTGCCGTCTGAATTTGACCTGAGCGCATTTTTACGC<br>GCCGGAGAAAACCGCCTCGCGGTGATGGTGCTGCGCTGGAGTGACGGCAGTTATCTGGAA<br>GATCAGGATATGTGGCGGATGAGCGGCATTTTCCGTGACGTCTCGTTGCTGCATAAACCG<br>ACTACACAAATCAGCGATTTCCATGTTGCCACTCGCTTTAATGATGATTTCAGCCGCGCTG<br>TACTGGAGGCTGAAGTTCAGATGTGCGGCGAGTTGCGTGACTACCTACGGGTAACAGTTT<br>CTTTATGGCAGGGTGAAACGCAGGTCGCCAGCGGCACCGCGCCTTTCGGCGGTGAAATTA<br>TCGATGAGCGTGGTGGTTATGCCGATCGCGTCACACTACGTCTGAACGTCGAAAACCCGA<br>AACTGTGGAGCGCCGAAATCCCGAATCTCTATCGTGCGGTGGTTGAACTGCACACCGCCG<br>ACGGCACGCTGATTGAAGCAGAAGCCTGCGATGTCGGTTTCCGCGAGGTGCGGATTGAAA<br>ATGGTCTGCTGCTGCTGAACGGCAAGCCGTTGCTGATTCGAGGCGTTAACCGTCACGAGC<br>ATCATCCTCTGCATGGTCAGGTCATGGATGAGCAGACGATGGTGCAGGATATCCTGCTGA<br>TGAAGCAGAACAACTTTAACGCCGTGCGCTGTTCGCATTATCCGAACCATCCGCTGTGGT<br>ACACGCTGTGCGACCGCTACGGCCTGTATGTGGTGGATGAAGCCAATATTGAAACCCACG<br>GCATGGTGCCAATGAATCGTCTGACCGATGATCCGCGCTGGCTACCGGCGATGAGCGAAC<br>GCGTAACGCGAATGGTGCAGCGCGATCGTAATCACCCGAGTGTGATCATCTGGTCGCTGG<br>GGAATGAATCAGGCCACGGCGCTAATCACGACGCGCTGTATCGCTGGATCAAATCTGTCG<br>ATCCTTCCCGCCCGGTGCAGTATGAAGGCGGCGGAGCCGACACCACGGCCACCGATATTA<br>TTTGCCCGATGTACGCGCGCGTGGATGAAGACCAGCCCTTCCCGGCTGTGCCGAAATGGT<br>CCATCAAAAAATGGCTTTCGCTACCTGGAGAGACGCGCCCGCTGATCCTTTGCGAATACG<br>CCCACGCGATGGGTAACAGTCTTGGCGGTTTCGCTAAATACTGGCAGGCGTTTCGTCAGT<br>ATCCCCGTTTACAGGGCGGCTTCGTCTGGGACTGGGTGGATCAGTCGCTGATTAAATATG<br>ATGAAAACGGCAACCCGTGGTCGGCTTACGGCGGTGATTTTGGCGATACGCCGAACGATC<br>GCCAGTTCTGTATGAACGGTCTGGTCTTTGCCGACCGCACGCCGCATCCAGCGCTGACGG<br>AAGCAAAACACCAGCAGCAGTTTTTCCAGTTCCGTTTATCCGGGCAAACCATCGAAGTGA |

| | |
|---|---|
| | CCAGCGAATACCTGTTCCGTCATAGCGATAACGAGCTCCTGCACTGGATGGTGGCGCTGG<br>ATGGTAAGCCGCTGGCAAGCGGTGAAGTGCCTCTGGATGTCGCTCCACAAGGTAAACAGT<br>TGATTGAACTGCCTGAACTACCGCAGCCGGAGAGCGCCGGGCAACTCTGGCTCACAGTAC<br>GCGTAGTGCAACCGAACGCGACCGCATGGTCAGAAGCCGGGCACATCAGCGCCTGGCAG<br>CAGTGGCGTCTGGCGGAAAACCTCAGTGTGACGCTCCCCGCCGCGTCCCACGCCATCCCG<br>CATCTGACCACCAGCGAAATGGATTTTTGCATCGAGCTGGGTAATAAGCGTTGGCAATTT<br>AACCGCCAGTCAGGCTTTCTTTCACAGATGTGGATTGGCGATAAAAAACAACTGCTGACG<br>CCGCTGCGCGATCAGTTCACCCGTGCACCGCTGGATAACGACATTGGCGTAAGTGAAGCG<br>ACCCGCATTGACCCTAACGCCTGGGTCGAACGCTGGAAGGCGGCGGGCCATTACCAGGCC<br>GAAGCAGCGTTGTTGCAGTGCACGGCAGATACACTTGCTGATGCGGTGCTGATTACGACC<br>GCTCACGCGTGGCAGCATCAGGGGAAAACCTTATTTATCAGCCGGAAAACCTACCGGATT<br>GATGGTAGTGGTCAAATGGCGATTACCGTTGATGTTGAAGTGGCGAGCGATACACCGCAT<br>CCGGCGCGGATTGGCCTGAACTGCCAGCTGGCGCAGGTAGCAGAGCGGGTAAACTGGCTC<br>GGATTAGGGCCGCAAGAAAACTATCCCGACCGCCTTACTGCCGCCTGTTTTGACCGCTGG<br>GATCTGCCATTGTCAGACATGTATACCCCGTACGTCTTCCCGAGCGAAAACGGTCTGCGCT<br>GCGGGACGCGCGAATTGAATTATGGCCCACACCAGTGGCGCGGCGACTTCCAGTTCAACA<br>TCAGCCGCTACAGTCAACAGCAACTGATGGAAACCAGCCATCGCCATCTGCTGCACGCGG<br>AAGAAGGCACATGGCTGAATATCGACGGTTTCCATATGGGGATTGGTGGCGACGACTCCT<br>GGAGCCCGTCAGTATCGGCGGAATTCCAGCTGAGCGCCGGTCGCTACCATTACCAGTTGG<br>TCTGGTGTCAAAAATAATTCAGCCAAAAAACTTAAGACCGCCGGTCTTGTCCACTACCTTG<br>CAGTAATGCGGTGGACAGGATCGGCGGTTTTCTTTTCTCTTCTCAA |
| pUCGA-trH – plasmid encoding T7 expression of trH RNA trigger | GGAAGTTTGTCTAGATCTCAGGCGTGGATGCGGGTACCGAGCTCGAATTCACTGGCCGTC<br>GTTTTACAACGTCGTGACTGGGAAAACCCTGGCGTTACCCAACTTAATCGCCTTGCAGCAC<br>ATCCCCCTTTCGCCAGCTGGCGTAATAGCGAAGAGGCCCGCACCGATCGCCCTTCCCAAC<br>AGTTGCGCAGCCTGAATGGCGAATGGCGCCTGATGCGGTATTTTCTCCTTACGCATCTGTG<br>CGGTATTTCACACCGCATATGGTGCACTCTCAGTACAATCTGCTCTGATGCCGCATAGTTA<br>AGCCAGCCCCGACACCCGCCAACACCCGCTGACGCGCCCTGACGGGCTTGTCTGCTCCCG<br>GCATCCGCTTACAGACAAGCTGTGACCGTCTCCGGGAGCTGCATGTGTCAGAGGTTTTCA<br>CCGTCATCACCGAAACGCGCGAGACGAAAGGGCCTCGTGATACGCCTATTTTTATAGGTT<br>AATGTCATGATAATAATGGTTTCTTAGACGTCAGGTGGCACTTTTCGGGGAAATGTGCGC<br>GGAACCCCTATTTGTTTATTTTTCTAAATACATTCAAATATGTATCCGCTCATGAGACAAT<br>AACCCTGATAAATGCTTCAATAATATTGAAAAAGGAAGAGTATGAGTATTCAACATTTCC<br>GTGTCGCCCTTATTCCCTTTTTTGCGGCATTTTGCCTTCCTGTTTTTGCTCACCCAGAAACG<br>CTGGTGAAAGTAAAAGATGCTGAAGATCAGTTGGGTGCACGAGTGGGTTACATCGAACTG<br>GATCTCAACAGCGGTAAGATCCTTGAGAGTTTTCGCCCCGAAGAACGTTTTCCAATGATG<br>AGCACTTTTAAAGTTCTGCTATGTGGCGCGGTATTATCCCGTATTGACGCCGGGCAAGAGC<br>AACTCGGTCGCCGCATACACTATTCTCAGAATGACTTGGTTGAGTACTCACCAGTCACAG<br>AAAAGCATCTTACGGATGGCATGACAGTAAGAGAATTATGCAGTGCTGCCATAACCATGA<br>GTGATAACACTGCGGCCAACTTACTTCTGACAACGATCGGAGGACCGAAGGAGCTAACCG<br>CTTTTTTGCACAACATGGGGGATCATGTAACTCGCCTTGATCGTTGGGAACCGGAGCTGA<br>ATGAAGCCATACCAAACGACGAGCGTGACACCACGATGCCTGTAGCAATGGCAACAACG<br>TTGCGCAAACTATTAACTGGCGAACTACTTACTCTAGCTTCCCGGCAACAATTAATAGACT<br>GGATGGAGGCGGATAAAGTTGCAGGACCACTTCTGCGCTCGGCCCTTCCGGCTGGCTGGT<br>TTATTGCTGATAAATCTGGAGCCGGTGAGCGTGGGTCTCGCGGTATCATTGCAGCACTGG<br>GGCCAGATGGTAAGCCCTCCCGTATCGTAGTTATCTACACGACGGGGAGTCAGGCAACTA<br>TGGATGAACGAAATAGACAGATCGCTGAGATAGGTGCCTCACTGATTAAGCATTGGTAAC<br>TGTCAGACCAAGTTTACTCATATATACTTTAGATTGATTTAAAACTTCATTTTTAATTTAAA<br>AGGATCTAGGTGAAGATCCTTTTTGATAATCTCATGACCAAAATCCCTTAACGTGAGTTTT<br>CGTTCCACTGAGCGTCAGACCCCGTAGAAAAGATCAAAGGATCTTCTTGAGATCCTTTTTT<br>TCTGCGCGTAATCTGCTGCTTGCAAACAAAAAAACCACCGCTACCAGCGGTGGTTTGTTTG<br>CCGGATCAAGAGCTACCAACTCTTTTTCCGAAGGTAACTGGCTTCAGCAGAGCGCAGATA<br>CCAAATACTGTTCTTCTAGTGTAGCCGTAGTTAGGCCACCACTTCAAGAACTCTGTAGCAC<br>CGCCTACATACCTCGCTCTGCTAATCCTGTTACCAGTGGCTGCTGCCAGTGGCGATAAGTC<br>GTGTCTTACCGGGTTGGACTCAAGACGATAGTTACCGGATAAGGCGCAGCGGTCGGGCTG<br>AACGGGGGGTTCGTGCACACAGCCCAGCTTGGAGCGAACGACCTACACCGAACTGAGAT<br>ACCTACAGCGTGAGCTATGAGAAAGCGCCACGCTTCCCGAAGGGAGAAAGGCGGACAGG<br>TATCCGGTAAGCGGCAGGGTCGGAACAGGAGAGCGCACGAGGGAGCTTCCAGGGGGAAA<br>CGCCTGGTATCTTTATAGTCCTGTCGGGTTTCGCCACCTCTGACTTGAGCGTCGATTTTTGT<br>GATGCTCGTCAGGGGGGCGGAGCCTATGGAAAAACGCCAGCAACGCGGCCTTTTTACGGT<br>TCCTGGCCTTTTGCTGGCCTTTTGCTCACATGTTCTTTCCTGCGTTATCCCCTGATTCTGTG<br>GATAACCGTATTACCGCCTTTGAGTGAGCTGATACCGCTCGCCGCAGCCGAACGACCGAG<br>CGCAGCGAGTCAGTGAGCGAGGAAGCGGAAGAGCGCCCAATACGCAAACCGCCTCTCCC<br>CGCGCGTTGGCCGATTCATTAATGCAGCTGGCACGACAGGTTTCCCGACTGGAAAGCGGG<br>CAGTGAGCGCAACGCAATTAATGTGAGTTAGCTCACTCATTAGGCACCCCAGGCTTTACA<br>CTTTATGCTTCCGGCTCGTATGTTGTGTGGAATTGTGAGCGGATAACAATTTCACACAGGA<br>AACAGCTATGACCATGATTACGCCAAGCTTGCATGCCTGCAGGTCGACTCTAGAGGATCG<br>AGCACATCTCGTTCGCTATTCAGGGATTGGGAAACACAGAAAAAAGCCCGCACCTGACAG<br>TGCGGGCTTTTTTTTTCGACCAAAGGTAATACGACTCACTATAGGGACCGTGGACCGCATG<br>AGGTCCACGGTAAACATAACTATAACAAGCCTACAATTCATTCAAACTTCAGCCAAAAAA<br>CTTAAGACCGCCGGTCTTGTCCACTACCTTGCAGTAATGCGGTGGACAGGATCGGCGGTTT |

| | |
|---|---|
| | TCTTTTCTCTTCTCAACTCGGTACCAAATTCCAGAAAAGAGACGCTTTCGAGCGTCTTTTTTCGTTTTGGTCCGTCAGTTTCACCTGTTTTACGTAAAAACCCGCTTCGGCGGGTTTTTACTTTTGG |
| mslp151 – forward primer 1 Colicin E1-6xHis | CAACGGTTTCCCTCTAGAAATAATTTTGTTTAACTTTAAGAAGGAGATATACATATGGAAACCGCGGTAGCGTAC |
| mslp152 – forward primer 2 Colicin E1-6xHis | AGATCTCGATCCCGCGAAATTAATACGACTCACTATAGGGAGACCACAACGGTTTCCCTCTAGAAATAATTTTGTT |
| mslp153 – reverse primer 1 Colicin E1-6xHis | AGCAGCCAACTCAGCTTCCTTTCGGGCTTTGTTAGCAGCCGGTCGACTTAATGATGATGATGATGATGAATCCCTAACACCTCATTTAT |
| mslp154 – reverse primer 2 Colicin E1-6xHis | CAAAAAACCCCTCAAGACCCGTTTAGAGGCCCCAAGGGGTTATGCTAGTTATTGCTCAGCGGTGGCAGCAGCCAACTCAGCTTCCTTTCG |

**Table S3** Final concentrations of components in three different CFPS buffers

| Component | PANOx-SP final concentration |
|---|---|
| Magnesium glutamate | 12 mM |
| Potassium glutamate | 130 mM |
| Ammonium glutamate | 10 mM |
| Magnesium acetate | 3.7 mM* |
| Potassium acetate | 16.02 mM* |
| Tris acetate | 2.67 mM (pH 8.2)* |
| HEPES | 57 mM (pH 7.4) |
| ATP | 1.2 mM |
| GTP | 0.85 mM |
| CTP | 0.85 mM |
| UTP | 0.85 mM |
| Folinic acid | 0.072 mM |
| tRNA | 170.6 μg/mL |
| Alanine | 2 mM |
| Arginine | 2 mM |
| Histidine | 2 mM |
| Lysine (monoHCl) | 2 mM |
| Aspartic acid | 2 mM |
| Glutamic acid | 2 mM |
| Isoleucine | 2 mM |
| Leucine | 2 mM |
| Methionine | 2 mM |
| Phenylalanine | 2 mM |
| Tryptophan | 2 mM |
| Tyrosine | 2 mM |

| | |
|---|---|
| Valine | 2 mM |
| Serine | 2 mM |
| Threonine | 2 mM |
| Asparagine | 2 mM |
| Glutamine | 2 mM |
| Cysteine | 2 mM |
| Glycine | 2 mM |
| Proline | 2 mM |
| PEP | 33 mM |
| NAD | 0.33 mM |
| CoA | 0.27 mM |
| Spermidine | 1.5 mM |
| Putrescine | 1 mM |
| Oxalic acid | 4 mM |
| T7 RNA polymerase | 100 µg/mL |
| Plasmid DNA | 6.4 nM |
| RNase Inhibitor | 0.8 U/µL |
| Cell extract | 26.7% v/v* |
| DTT | 0.8 mM* |

* Items added as part of the cell extract solution

**Table S4.** List of supplementary videos

| File | Description |
|---|---|
| Video S1_2H No DNA | Tiled microscopy video of whole CFPS-PLGA film **with no DNA** added. DIC, green (sfGFP), and red (mRuby) fluorescence channels are merged. |
| Video S2_2H sfGFP DNA | Tiled microscopy video of whole CFPS-PLGA film with **sfGFP DNA** added. DIC, green (sfGFP), and red (mRuby) fluorescence channels are merged. |
| Video S3_2H mRuby DNA | Tiled microscopy video of whole CFPS-PLGA film with **mRuby DNA** added. DIC, green (sfGFP), and red (mRuby) fluorescence channels are merged. |
| Video S4_2H both DNA | Tiled microscopy video of whole CFPS-PLGA film with **both sfGFP and mRuby DNA** added. DIC, green (sfGFP), and red (mRuby) fluorescence channels are merged. |
